# Centromeric transposable elements and epigenetic status drive karyotypic variation in the eastern hoolock gibbon

**DOI:** 10.1101/2024.08.29.610280

**Authors:** Gabrielle A. Hartley, Mariam Okhovat, Savannah J. Hoyt, Emily Fuller, Nicole Pauloski, Nicolas Alexandre, Ivan Alexandrov, Ryan Drennan, Danilo Dubocanin, David M. Gilbert, Yizi Mao, Christine McCann, Shane Neph, Fedor Ryabov, Takayo Sasaki, Jessica M. Storer, Derek Svendsen, William Troy, Jackson Wells, Leighton Core, Andrew Stergachis, Lucia Carbone, Rachel J. O’Neill

## Abstract

Great apes have maintained a stable karyotype with few large-scale rearrangements; in contrast, gibbons have undergone a high rate of chromosomal rearrangements coincident with rapid centromere turnover. Here we characterize assembled centromeres in the Eastern hoolock gibbon, *Hoolock leuconedys* (HLE), finding a diverse group of transposable elements (TEs) that differ from the canonical alpha satellites found across centromeres of other apes. We find that HLE centromeres contain a CpG methylation centromere dip region, providing evidence this epigenetic feature is conserved in the absence of satellite arrays; nevertheless, we report a variety of atypical centromeric features, including protein-coding genes and mismatched replication timing. Further, large structural variations define HLE centromeres and distinguish them from other gibbons. Combined with differentially methylated TEs, topologically associated domain boundaries, and segmental duplications at chromosomal breakpoints, we propose that a “perfect storm” of multiple genomic attributes with propensities for chromosome instability shaped gibbon centromere evolution.

## Introduction

Gibbons (Family Hylobatidae) are a group of ∼20 species of small apes that last shared a common ancestor with their closest relatives, great apes, ∼17 million years ago. As a consequence of high rates of inter- and intra-chromosomal rearrangements, the karyotypes of the four extant gibbon genera are highly diverse with variable chromosome numbers (Figure 1A, 1B)^1–3^: *Hoolock*, (2n=38); *Nomascus* (2n=52); *Symphalangus* (2n=50) and *Hylobates* (2n=44)^4,5^. In gibbons, rapid chromosomal rearrangements have led to a high rate of centromere turnover; for example, the eastern hoolock gibbon (*Hoolock leuconedys*, HLE) retains over 50 chromosome rearrangements leading to the inactivation of 13 centromeres and the formation of six evolutionary new centromeres (ENCs)^2^. Moreover, unlike the centromeres of great apes that are composed of 171 bp AT-rich alpha satellite arrays and larger higher order arrays (HORs)^6–8^, the centromeres of many gibbon species lack large alpha satellite arrays arranged in an HOR structure^9,10^.

**Figure 1:**
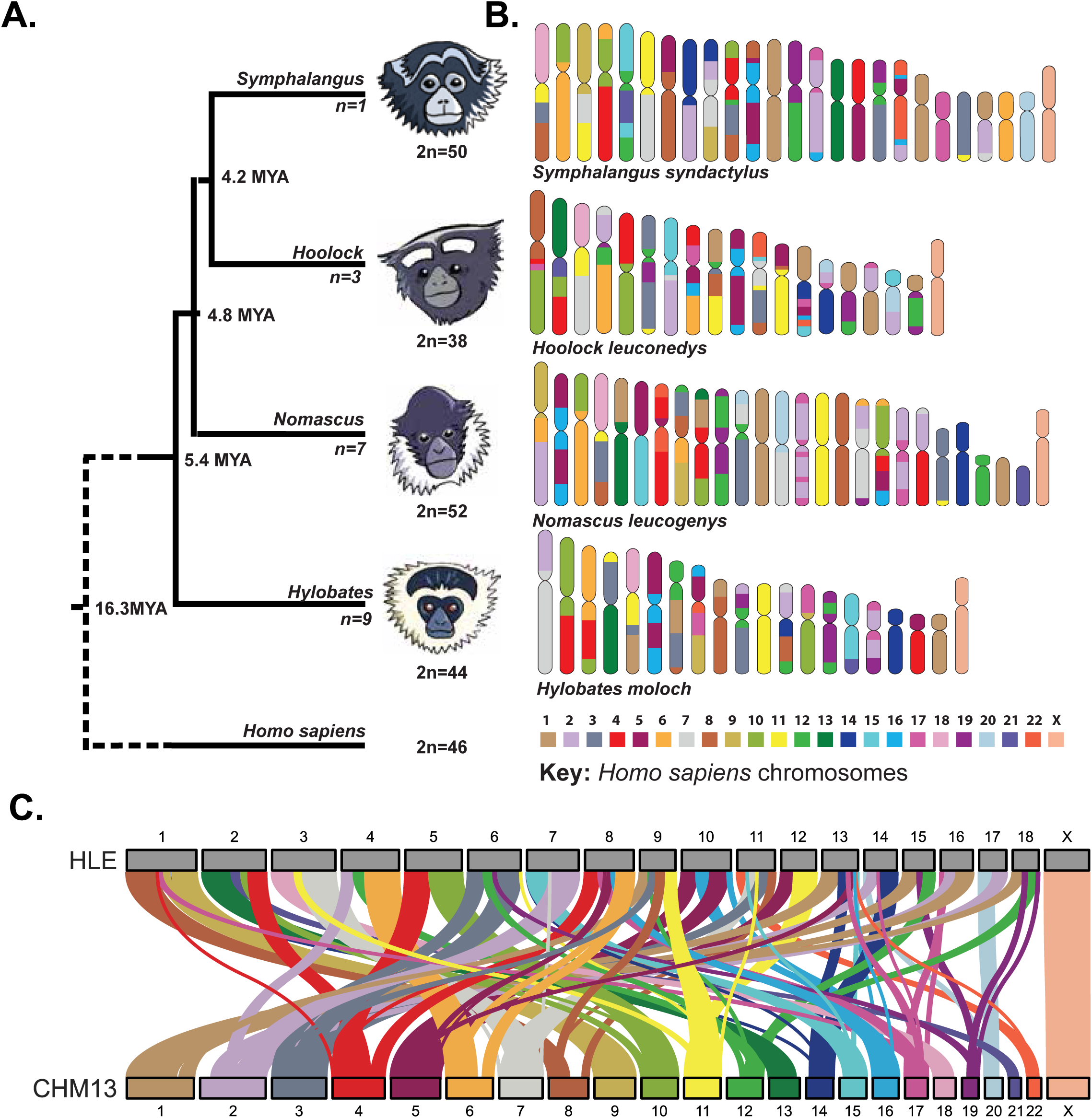
Gibbon genera display high rates of karyotype variation since their radiation ∼8 million years ago. **(A)** The phylogeny of lesser apes with estimated divergence times is depicted based on ^81^. The four gibbon genera (*Hylobates*, *Hoolock*, *Symphalangus*, and *Nomascus*) descended from a shared common ancestor ∼8 million years ago and now present with highly derivative karyotypes (ranging from 2n=38 to 52). The number below each branch represents the number of known species within each genus. **(B)** Synteny between human and gibbon chromosomes is shown with a representative species from each gibbon genus, based on ^2^. Each color represents homology to a different human autosome, with a key depicted below. **(C)** Synteny between our assembled HLE chromosomes (top) and human T2T-CHM13 chromosomes (bottom) agree with those demonstrated in **(B)**, confirming lack of large-scale structural mis-assemblies and highlights the genome-wide chromosome rearrangements present in the HLE genome.

Recent studies in a complete human genome have identified a variety of epigenetic features that define centromere function, including a dip of decreased CpG methylation at the active centromere (centromere dip region, CDR)^11,12^, late replication timing^13^, and variable chromatin compaction into dichromatin^14^. However, the conservation and function of these features beyond humans and within satellite-free centromeres is yet to be resolved. In addition, despite gibbons exhibiting a rate of chromosomal rearrangements up to 20 times higher than other primates^2^, the factors driving this high rate are still not fully understood. One potential contributor to karyotype variation in gibbons is the propagation of LAVA, an active ∼2 kilobase gibbon-specific composite retrotransposon composed of **L**INE, ***A****lu*Sz, variable number tandem repeat (**V**NTR), and ***A****lu*-like segments^9,15^. While present amongst all gibbons, LAVA has propagated at variable rates among the genera and are most abundantly found in the centromeres and pericentromeres of *Hoolock* species^9^. LAVA elements are hypothesized to contribute to karyotype evolution by their subsequent co-option as a regulatory element within genome repair pathways^16^. Although it is unclear whether LAVA arrays and centromeric variation are a causative or consequential factor in gibbon speciation, these highly variable centromeric units are a distinguishing feature of the *Hoolock* genus. Thus, the eastern hoolock gibbon serves as a compelling model to interrogate the potential relationship between rearrangement breakpoints and centromere turnover during rapid karyotypic evolution.

To survey centromeres and chromosomal rearrangements of the eastern hoolock gibbon, we developed a long-read based genome assembly for HLE, *cmHooLeu*1. By analyzing the centromere-specific histone centromere protein A (CENP-A) occupancy, transposable elements, CpG methylation, chromatin accessibility, replication timing, transcription, RNA polymerase occupancy, and genome spatial conformation, we identified features that define centromere identity and breakpoints in the HLE genome. We find that, despite variability in transposable element content and sequence identity, HLE centromeres have a CDR that overlaps with CENP-A enrichment, indicating this epigenetic feature is not restricted to alpha satellite-containing centromeres. We further identify genomic and epigenomic features within functional centromeres associated with replication stress and chromosome instability, such as the presence of protein-coding genes, regulatory elements, pericentromeric segmental duplications of LAVA and SST1 repeats, and variable replication timing within the centromere. Thus, we hypothesize that a combination of genomic and epigenetic features combine to create a “perfect storm” associated with an increased propensity for chromosomal rearrangements.

## Results

### Generating and annotating a genome assembly for Betty, a female eastern hoolock gibbon

To assemble a reference genome for Betty, a female HLE, we generated ∼166 Gb of DNA sequence across PromethION flow cells from Oxford Nanopore Technologies (∼59x coverage, including 59 Gb of ONT ultra-long sequencing), 62x coverage of Illumina PCR-free sequencing, and 29x coverage of Dovetail Omni-C sequencing from a lymphoblastoid cell line. Following assembly, scaffolding, and polishing, the final reference, *cmHooLeu*1, consists of 19 scaffolds corresponding to the haploid chromosome number of HLE with a total length of 2.761 Gb (N50 = 159.997 Mb), roughly equivalent to the short-read *k*-mer based genome estimate of 2.779 - 2.782 Gb (Table 1). Alignment to the human reference genome T2T-CHM13 confirms the organization of HLE to human syntenic blocks along the HLE chromosomes^2,11^, supporting the lack of large structural mis-assemblies in our HLE assembly (Figure 1C). Using the Benchmarking Universal Single-Copy Ortholog (*BUSCO*) analysis ^17^, the expected gene set is 95.5% complete using the primates dataset and the QV score using Illumina PCR-free sequencing was determined to be 46.71, roughly equivalent to an inferred nucleotide accuracy of 99.997% (Table 1)^18^.

**Table 1:** HLE assembly quality statistics. Basic quality statistics pertaining to contiguity, length, GC percentage, and number of Ns were defined by *QUAST*. Gene completeness was estimated using the primates odb10 BUSCO database, and *k*-mer completeness was estimated using *Merqury*.

|  |  |
| --- | --- |
| Number of scaffolds | 19 |
| **Total length (bp)** | 2,760,609,521 |
| **Largest contig (bp)** | 226,217,378 |
| **GC (%)** | 41.25 |
| **N50** | 159,996,863 |
| **N75** | 115,684,913 |
| **L50** | 8 |
| **L75** | 13 |
| **# N's per 100 kbp** | 0.22 |
| **BUSCO complete** | 95.5% |
| **BUSCO fragmented** | 1.2% |
| **BUSCO missing** | 3.3% |
| **Merqury QV** | 46.71 |
| **Merqury Completeness** | 97.87 |
| **Number of protein coding genes** | 20,113 |

Using RepeatMasker^19^, we identified an overall repeat content of 50.38%, for a total of 1,390,689,114 repetitive bases composed primarily of LINEs (21.54%), SINEs (14.09%), and LTRs (8.57%) (Table S1). LAVA comprises 0.7% of the assembled genome, higher than found in genomes of other gibbon genera, (0.2% in *Symphalangus syndactylus*, GCF_028878055.2; 0.12% in *Nomascus leucogenys,* GCF_006542625.1; and 0.06% *Hylobates moloch,* GCF_009828535.3). A total of 20,113 protein-coding genes were predicted (BUSCO: 99.5% complete; 0.2% fragmented; 0.3% missing; n=9226). In addition, to assess the nascent transcriptome, we performed precision run-on sequencing (PRO-seq)^20^. Regardless of the alignment method, the top three transcriptionally active repeats are SINEs, LINEs, and LTRs, and satellite transcription was low, mimicking the pattern seen in humans (Figure S1, S2)^21^.

To identify centromeric regions within the HLE assembly that nucleate kinetochore assembly initiated through the deposition of CENP-A, CUT&RUN sequencing was performed using an antibody for centromere protein A (CENP-A). One prospective centromere region was identified per chromosome based on elevated CENP-A CUT&RUN read coverage. Six centromeres (Cen1, Cen3, Cen8, Cen9, Cen11, and CenX) displayed generally even ONT read coverage across the region, indicating uncollapsed assembly of these centromeric sequences (Figures S3, S4, Table S2), and thus, unless noted otherwise, only these six centromeres were used in subsequent analyses.

### HLE centromeres vary in repeat organization, yet maintain a centromere dip region (CDR) and a dichromatin conformation

Among assembled centromeres, the span of CENP-A enrichment averaged 130 kilobases (75-162 kb) (Table S2), nearly half the average size of CENP-A enrichment in CHM13 centromeres (317 kb)^8^. Unlike the alpha satellite-rich centromeres of great apes, HLE centromeres displayed highly variable, complex repeat content dominated by the presence of transposable elements (TEs) both at the surrounding pericentromere (Figure 2A) and site of CENP-A enrichment (Figure 2B). The most prevalent TEs across assembled centromeres were long interspersed nuclear elements (LINEs), with presence in four of the six uncollapsed centromeres (Cen1, Cen3, Cen8, and Cen9), followed by a mixture of short interspersed nuclear elements (SINEs) and long terminal repeats (LTRs) (Figure 2B, Table S3). Notably, only two of the centromeres, Cen3 and CenX, were composed of the primate centromeric repeat alpha satellite (Figure 2A, 3B, S5B, S5D, Table S3), and only in CenX was it the major constituent of the CENP-A enriched region (Figure 2B, 3A). Other satellites were also identified in pericentromeric regions; namely Cen3 contained centromeric repeats (CER), a 48 bp repeat found on the centromeric q arms of human chromosomes 22, 14, and on chromosome 18^22^; Cen8 contained beta satellites (BSR)^23,24^, which are enriched in heterochromatin in human chromosomes; and Cen8 and CenX contained gamma satellites (GSAT), a GC-rich satellite with large arrays in humans and often associated with alpha satellite DNA (Figure 2A, Table S3)^25^. While LAVA was not found to be the main constituent of the DNA associated with CENP-A, it was found to be present in three of the analyzed centromeres (Cen8, Cen9, and Cen11).

**Figure 2:**
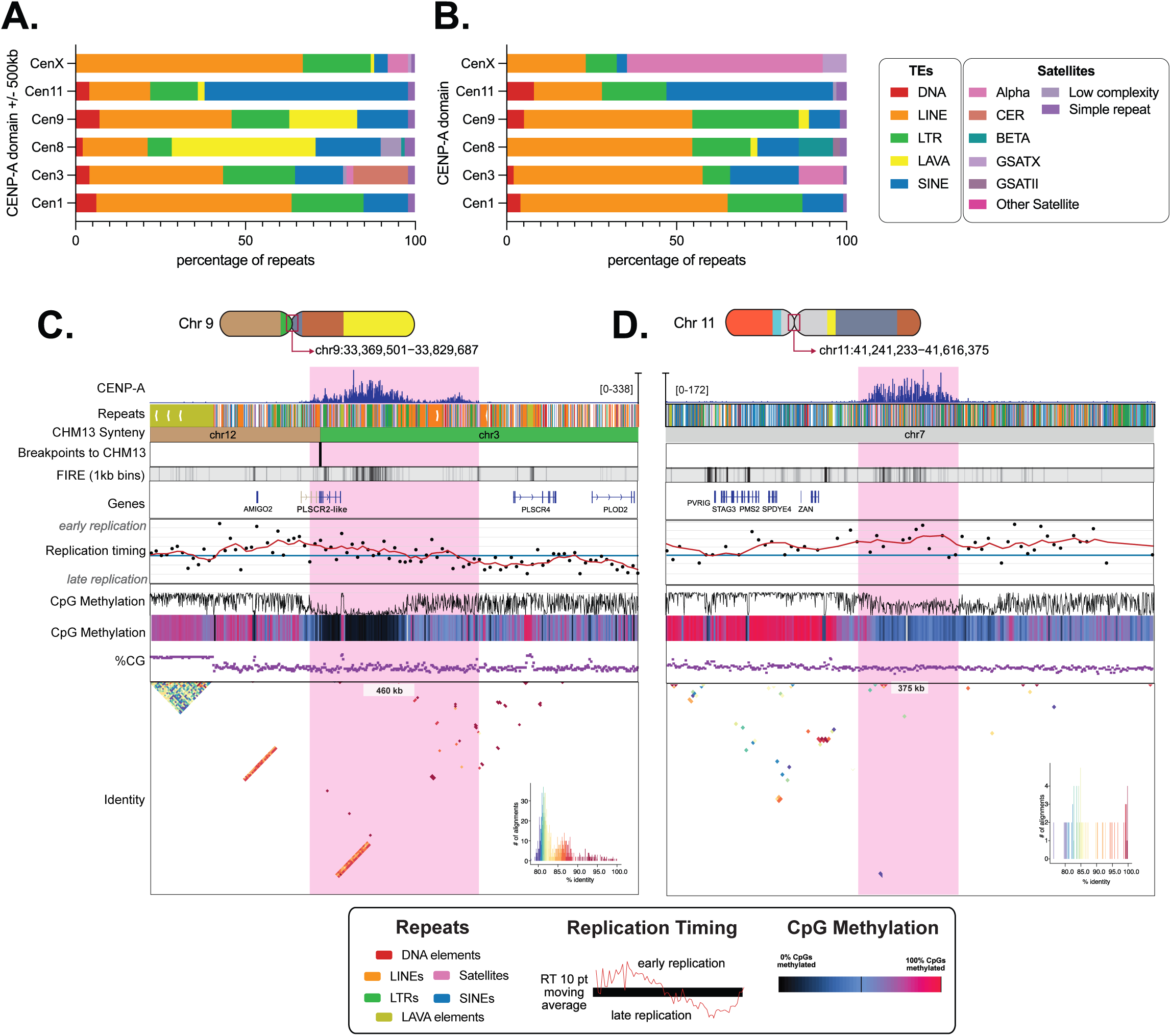
Centromeres of the Eastern hoolock gibbon are enriched with diverse transposable elements and vary in repeat organization, yet maintain a CDR. The percentage of total repeat content (bp) classified as each repeat class is shown for **(A)** the centromere region (defined as the CENP-A domain and 500kb upstream and downstream) and **(B)** for the CENP-A enrichment domain of the six assembled HLE centromeres, highlighting the highly variable repeat composition of centromeres. Below, chromosome ideograms show the position of HLE Cen9 **(C)** and Cen11 **(D)** with chromosomes colored by their synteny to human chromosomes as per Figure 1. From top to bottom, genome tracks denote CENP-A CUT&RUN enrichment (blue), repeat annotations colored according to the key below, synteny to T2T-CHM13, and predicted HLE-CHM13 synteny breakpoints. Fiber-seq inferred regulatory element (FIRE) tracks show FIRE density binned per 1kb on a heatmap scale from white to black (i.e. low to high density), showing increased density of FIREs correlating with CENP-A enrichment corresponding to dichromatin organization. Gene tracks (blue and tan bars indicating true and falsely predicted exons, respectively) show gene predictions from FLAG, showing the presence of several genes nearby and overlapping with CENP-A enrichment. Replication timing from E/L Repli-seq is shown as black points indicating the log ratio of early-to-late coverage over 5kb windows from 4 (early replication) to −4 (late replication), with a red line indicating the 10 point moving average. CpG methylation is shown via line plot (black line) and on a heatmap scale from low CpG methylation (black) to high CpG methylation (red). In HLE, CENP-A enrichment is associated with a dip in CpG methylation (CDRs) even in the absence of alpha satellite-containing centromeres and despite significant changes in CG density (purple). Finally, sequence identity plots are shown for each assembled centromere, with a scale from blue (low identity) to red (high identity). Overall, regions of CENP-A enrichment share little sequence identity compared to canonical and pericentromeric primate alpha satellite arrays.

**Figure 3:**
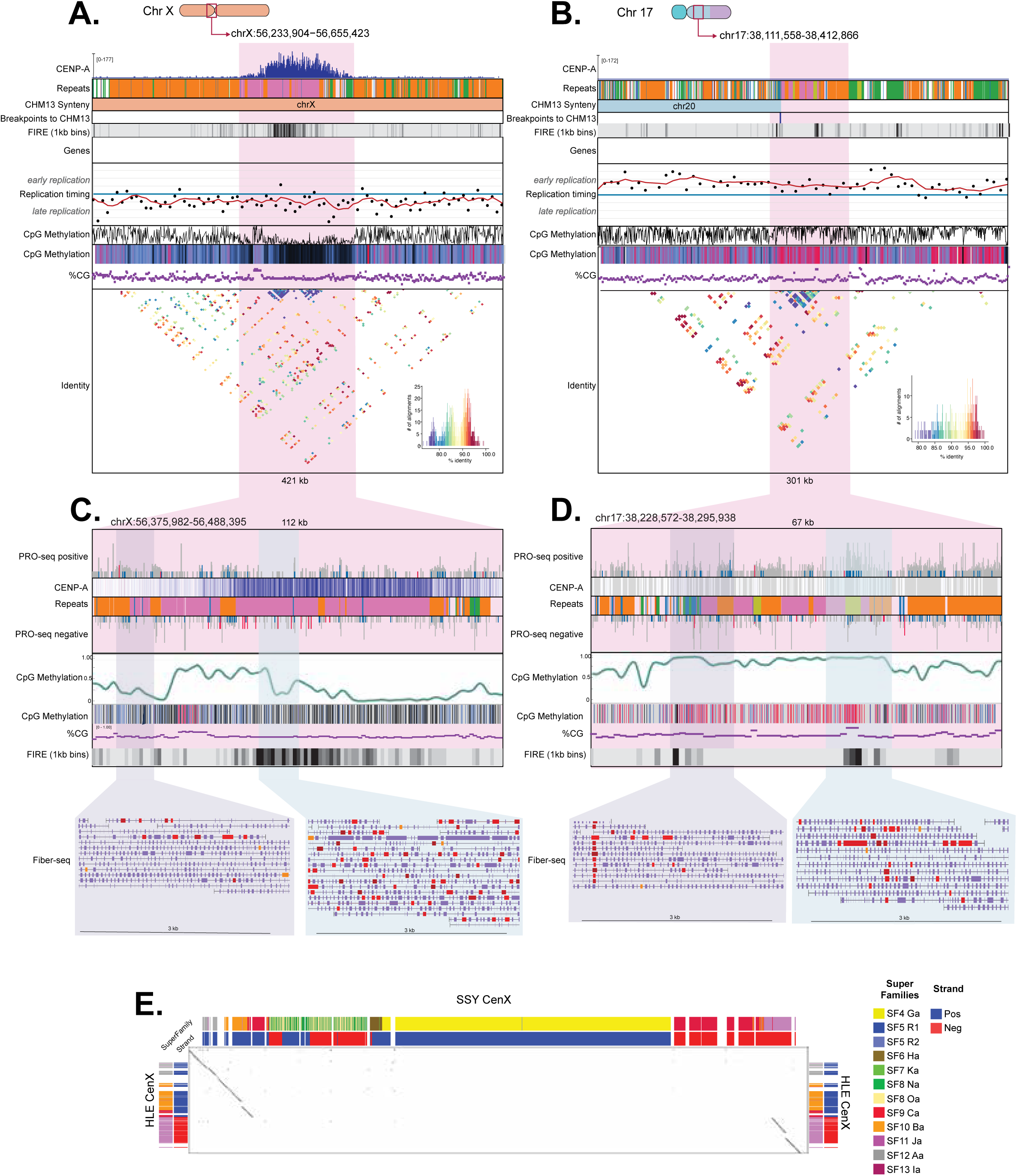
A latent alpha satellite centromere on HLE chromosome 17 lost epigenetic signatures of centromere function. Chromosome ideograms show the position of HLE CenX **(A)** flanked by dense LINE-rich regions and a latent centromere on Chr 17 **(B)** with chromosomes colored by their synteny to human chromosomes as per Figure 1. Genome tracks denote CENP-A CUT&RUN enrichment (blue), repeat annotations with each repeat class represented by a different color, synteny to T2T-CHM13, predicted breakpoints, FIRE elements, genes, replication timing, CpG methylation, CG percentage, and sequence identity, per the key in Figure 2. Zoomed panels for CenX **(C)** and Chr 17 **(D)** highlight the repeat organization of the two alpha satellite arrays, which present with LINE (CenX and Chr 17), *Alu* (CenX), and LAVA (Chr 17) insertions. Tracks show the presence of a CpG methylation dip region over the functional CenX, which is absent in the highly methylated latent alpha satellite centromere on Chr 17. Below, red boxes within FIRE tracks show the presence of disorganized FIRE elements/open chromatin (**C**, blue inset) in the active CenX corresponding to dichromatin, which is absent in the surrounding heterochromatin (**C**, purple inset) and the latent centromere on Chr 17 (**D**, purple inset). The LAVA element on Chr 17 (yellow) is more accessible than the surrounding alpha satellites (**D**, blue inset), suggesting it is functional. **(E)** The panel shows a dot-plot (Gepard, word length 50)^82^ comparing the HLE CenX (HLE_Chr_X:56370779-56477597) sequenced assembled herein to the SSY gibbon CenX (mSymSyn1_v2.0 chrX_hap1:70,696,431-71,324,358) described previously^27^. Corresponding alpha satellite annotation tracks are shown for both centromeres, showing alpha satellite super families (SFs) and the strand orientation (blue and red). A deletion breakpoint coincides with the only AS strand switch point in the HLE CenX, and is shown in the HLE_Chr_X:56442082-56442094 window. Breaks in the diagonals on both sides represent small deletions in SSY relative to the HLE-SSY common ancestor. UCSC Browser annotation tracks are described in^27^ and represent alpha satellite super family annotation (upper/left panels) and alpha satellite strand annotation (bottom/right panels).

All assembled centromeres lack the homogenous satellite higher order repeat (HOR) expansions concomitant with a highly identical core identified in humans and other primates (Figure 2C, 2D, 3A, S5)^8,26–28^. In fact, with the exception of alpha satellites sharing 75-80% sequence identity at the core of CenX, no other assembled centromere contains a highly homogenized (i.e. high sequence identity) repeat structure. Comparison to the published CenX of siamang (SSY)^27^ revealed that the SSY CenX is much larger (∼602 kb) compared to the HLE CenX (∼90 kb), and the two centromeres share only the extreme flanks of the alpha satellite array, with the middle, including the homogenous active HOR array in SSY, deleted in HLE CenX (Figure 3E). Analysis of alpha satellite suprachromosomal families (SFs) in HLE CenX array shows that it contains only the divergent monomeric layers formed by older SFs 9-12, which have not been shown to form functional centromeres in other apes. HLE CenX also does not contain SF4, which forms active centromeres in SSY^27^. Nevertheless, this reduced centromere is capable of supporting kinetochore formation.

Despite the lack of overall homogenous sequence identity at the centromere core in HLE, satellites arrays with high sequence identity repeats often were found to flank the site of CENP-A enrichment, including a CER satellite array on Cen3, a SINE/AluSp expansion on Cen8, a LAVA expansion on Cen9, and large tracts of LINE/L1s on CenX (Figure 2C, 2D, 3A, S5C). The variation in centromere composition suggests that gibbon centromeres underwent an independent evolutionary history compared to great apes, potentially as a response to drastic chromosomal rearrangements impacting centromeric regions. Like human alpha satellite centromeres^21^, these satellites exhibit low PRO-seq signal, yet PRO-seq signal increases over TEs that comprise the centromere and at the edges of CENP-A enrichment (Figure S6). However, unlike highly homogenized human centromere cores, the CENP-A binding domain is not a transcriptional dead zone, even at the alpha-satellite enriched CenX (Figure S6), mimicking the pattern seen at the diverged, outer alpha satellite layers in human centromeres^21^.

Within the active centromeric alpha satellite HORs in humans, a small region is associated with a distinctive decrease in CpG methylation (centromere dip region, CDR) coincident with CENP-A nucleosomes, indicating the importance of CpG methylation in proper kinetochore positioning and centromere functioning^12^. However, it is not clear if this epigenetic feature is conserved widely across centromeres and among small apes that lack the canonical ape centromeric alpha satellite organization. We took advantage of the single-molecule DNA modification detection obtainable with ONT sequencing reads to investigate CpG methylation across the assembled HLE centromeres and found that each contained a dip in methylation concomitant with CENP-A occupancy, despite high CpG methylation in the surrounding heterochromatin (Figure 2C, 2D, 3A, S5). The conservation of CDRs in HLE centromeres supports a model in which epigenetic regulation of kinetochore positioning is independent of alpha satellite-dense centromeres or HOR organization. Using Fiber-seq, we overlapped the six assembled HLE centromeres with m6A methylation calls to examine chromatin compaction in these regions and found a higher density of 6mA methylated regions and Fiber-seq inferred regulatory elements (FIREs) within CENP-A enrichment than outside, indicative of accessible chromatin (Figure 2C, 2D, 3A, S5). Thus, in addition to CDRs, all HLE centromeres contain “dichromatin”, a unique form of chromatin compaction recently reported to define centromeric chromatin^14^.

### Rearrangement of protein-coding genes are present throughout HLE centromeres and pericentromeres

Centromeres are typically established on highly condensed, gene-poor repetitive sequences, providing little opportunity for complex kinetochore machinery to interfere with the expression of protein coding genes^29^. However, we identified a total of 34 predicted genes within 200 kb of the span of CENP-A enrichment in the six assembled centromeres (Table S4, Figure 2C, 2D, 3A, S5). Of the annotated genes, GO biological processes with the highest enrichment combined scores included negative regulation of protein kinase activity by regulation of protein phosphorylation (GO:0044387, *CDK5RAP3*), response to DNA damage checkpoint signaling (GO:0072423, *EME1*), and antifungal innate immune response (GO:0061760, *DEFB119* and *DEFB4A*) (Table S5).

In two of the assembled centromeres, Cen8 and Cen9, predicted genes directly overlapped with CENP-A enrichment. Sequence analysis of the gene prediction overlapping with Cen8 revealed high (>80%) similarity to *CLUHP* pseudogenes; therefore, we more carefully annotated the Cen8 region by aligning human *CLUHP* exons to the region and found two full length *CLUHP* pseudogenes near HLE Cen8, which is also a breakpoint in HLE (Figure S5C, S7). In humans and in HLE Cen8, *CLUHP* pseudogenes flank segmental duplications and BSAT and LSAU-BSAT arrays, suggesting that these regions of high homology may have facilitated recombination events leading to chromosome rearrangements in HLE.

Gene annotation of Cen9 revealed that CENP-A enrichment overlapped a portion of a phospholipid scramblase 2 (*PLSCR2*) prediction, which is located in humans on a syntenic portion of chromosome 3 (Figure 2C, S5B). CENP-A enrichment at Cen9 overlaps with an HLE breakpoint between segments syntenic to human chromosome 3 and chromosome 12 and disrupts the phospholipid scramblase gene cluster present in humans on chromosome 3, comprised of *PLSCR1, PLSCR2, PLSCR4,* and *PLSCR5*, separating these genes to HLE chromosomes 6 (*PLSCR5, PLSCR1*) and 9 (*PLSCR4*) (Figure S8). While the HLE Cen9 region annotation is analogous to a potential *PLSCR2* transcript in humans, we found no evidence of *PLSCR2* exons at the HLE Cen9 region or elsewhere in the assembly, suggesting potential loss of the gene during recombination. In humans, *PLSCR2* is a lowly expressed protein-coding gene in a family of calcium binding proteins suggested to be involved in the blood coagulation cascade as well as macrophage clearance of apoptotic cells^30^. No evidence of transcription in HLE was detected in the recombined region by mapping total RNA-sequencing and precision nuclear run-on sequencing (PRO-seq) reads to the assembly (Figure S8). Despite the disruption to organization in HLE, *PLSCR1* remained highly transcribed in HLE and NLE gibbons (Figure S8).

### Segmental duplications and divergent replication timing, hallmarks of chromosome instability, are predicted among HLE centromeres

Highly identical and repetitive by definition, segmental duplications (SDs) are known to lead to chromosomal rearrangements via nonallelic homologous recombination, resulting in deletions, duplications, translocations, and inversions^31,32^. Notably, many studies have identified an association between SDs and subsequent chromosomal instability and recurrent genomic disorders^32–35^, and SDs have been linked to numerous cases of evolutionary chromosome rearrangements, including in mice^36^, chimpanzees^37^, and other apes^32,38,39^. Therefore, we decided to survey segmental duplications (SDs), broadly defined as >1kb long genomic duplications exceeding >90% sequence identity in the HLE assembly using *BISER*^35,40^. In total, 2,842 SDs were predicted, accounting for a total of 19,696,320 bp, or 0.71% of the assembly (Figure S9A). Intersecting segmental duplications with all centromere predictions ± 500kb, 10.6% of bases are predicted to be covered by SDs, nearly ∼15x higher than the genome-wide total. While the total predicted SDs are relatively low compared to SDs found in the human T2T-CHM13 genome (estimated to comprise 218 Mb and ∼7.0% of the genome^41^), we suspect collapsed sequences in our assembly result in an underestimation of SDs both genome-wide and among centromeres in our assembly.

Previous studies have reported associations between SST1 repeats, LAVA elements, and heterochromatic enrichment in HLE^9,10^. In our assembly, we found that CENP-A is not directly associated with LAVA nor SST1 enrichment, except for Cen6 and Cen18, respectively (Figure S4). Instead, SST1 and LAVA form large high identity arrays, sometimes exceeding 1Mb in size, in the pericentromere of many chromosomes (Figure 2C, 2D, S3, S4). Therefore, we extended our analysis of SDs to pericentromeric LAVA and SST1 repeats by intersecting SDs with LAVA and SST1 annotations. Of the 2,842 SDs, 154 (362,610 bp) contained LAVA (Figure S9B), while 487 SDs (91,899 bp) overlapped SST1s (Figure S9B). The majority of these high-identity SDs overlapped with pericentromeric locations (including both collapsed and uncollapsed centromeres), suggesting that pericentromeric loci are enriched in inter-chromosomal duplications (Figure S9B).

In addition to SDs, dysregulation of replication timing profiles has been linked extensively to genome instability, with observable shifts to earlier replication timing profiles associated with cancerous phenotypes^42–45^, chromosomal breakage^46,47^, and a higher probability of translocations in mice and humans^48^. In humans, the timing of centromere replication occurs during mid-late S phase and its exact timing is more precisely conserved across all chromosomes within an individual, suggesting the functional coordination of centromeric replication timing among chromosomes^13^. In order to investigate whether this applies to HLE centromeres that have variable transposable elements and gene composition, we performed E/L Repli-seq^49^ on HLE to map DNA replication timing genome-wide.

Among the six assembled HLE centromeres, three (Cen1, Cen3, and CenX) were consistent with expectations of mid to late S phase replication timing across the centromeric and pericentromeric regions (Figure 3A, S5A, S5B). However, Cen8, Cen9, and Cen11 showed evidence of mid to early S phase replication (Figure 2C, 2D, S5C). Additionally, while Cen11 appears to have consistent early S phase replication timing across the centromere region, Cen8 and Cen9 coincide with shifts in replication timing from early to late S phase, particularly near expansive LAVA arrays in the surrounding pericentromere (Figure 2C, S5C). Of note, the three assembled HLE centromeres with early S phase replication also share genes within or directly upstream of CENP-A enrichment, significant regions of upstream hypomethylation concomitant with increases in chromatin accessibility in the region, and pericentromeric LAVA expansions (Figure 2C, 2D, S5C), which may contribute to replication timing dysregulation.

### HLE Cen17 is defined by a unique composite repeat duplication not found in other apes

Within the *Hoolock* genus, HLE Cen17 and Cen11 (Figure 2D) are the only evolutionary new centromeres unique to the genus and they exhibit significantly lower heterochromatic LAVA amplification compared to all other centromeres^2,9^, prompting further investigation. Previous studies have identified a latent, non-functional centromere on the q arm of HLE chromosome 17 concomitant with the formation of an evolutionary new centromere (ENC)^2^. We therefore searched for alpha satellite sequences and identified a ∼28 kb array of alpha satellites presumed to be the latent centromere on chromosome 17, and corresponding to a portion of the syntenic centromere of SSY chromosome 24 (Figure S10). This array is smaller than most active alpha satellite arrays in primates, but *NucFreq*^50^ analysis supports proper assembly of the region, showing no evidence of read coverage anomalies nor assembly collapse (Figure S11). No CENP-A CUT&RUN reads aligned to the alpha satellite array located roughly ∼550 kb downstream of the CENP-A region (Figure 3B). Unlike active alpha satellite arrays in other primates^26^, this array lacks sequence homogenization and contains multiple transposable element insertions, including three LINE/L1Hylobs and two LAVA insertions, LAVA_B and LAVA_E (Figure 3B, 3D). CpG methylation analysis showed these satellites also lack a detectable CDR (Figure 3B, 3D), indicating alpha satellites arrays without CENP-A also lack a CDR. Fiber-seq analysis showed an absence of accessible chromatin compared to the active alpha satellite array on the X chromosome with the exception of a single highly accessible and potentially transposition competent LAVA insertion (Figure 3A, 3B). Further analyses are required to determine whether alpha satellite sequence degeneration, total array size reduction, and/or increased transposable element insertions into this region served as causative factors for centromere relocalization or followed centromere inactivation.

While Cen11 was successfully resolved and found to be composed of diverse TEs (Figure 2D), sequence collapses were present in the span of CENP-A enrichment of Cen17 (Figure S4). Nevertheless, we confirmed the span of CENP-A enrichment by overlapping mapped reads with unique 21-mer sequences in the HLE assembly localized to the expected 84 kb region (Figure 4A). Our analyses revealed that HLE Cen17 is composed of a composite repeat^21^ exclusive to this locus (Figure 4A). Each composite repeat contained 10 subunits: five LINEs (including one L2a and three L1Ms, one split by an *Alu* insertion) and five SINE/*Alu*s (including three *Alu*Sx elements, one *Alu*Jb, and one *Alu*Y), and a short (T)n simple repeat, averaging 3,319 bp in length (Figure 4B, Table S6) and arrayed in linear assembled sequence 24 times (Figure 4A). Fluorescence *in situ* hybridization (FISH) confirmed the centromeric localization of the sequence, hereafter called “L5A5”, to one chromosome pair confirmed to be HLE chromosome 17 (Table S7, Figure 4C). Although 24 copies of the L5A5 repeat were assembled in Cen17 (Figure 4D), we employed a *k*-mer based approach to estimate the uncollapsed copy number of the L5A5 repeat using Illumina PCR-Free WGS reads from the same individual^11^. Using this approach, we estimate 1,154 total copies of L5A5 genome-wide, or ∼577 copies per haplotype (∼1.9 Mb; Figure 4D).

**Figure 4:**
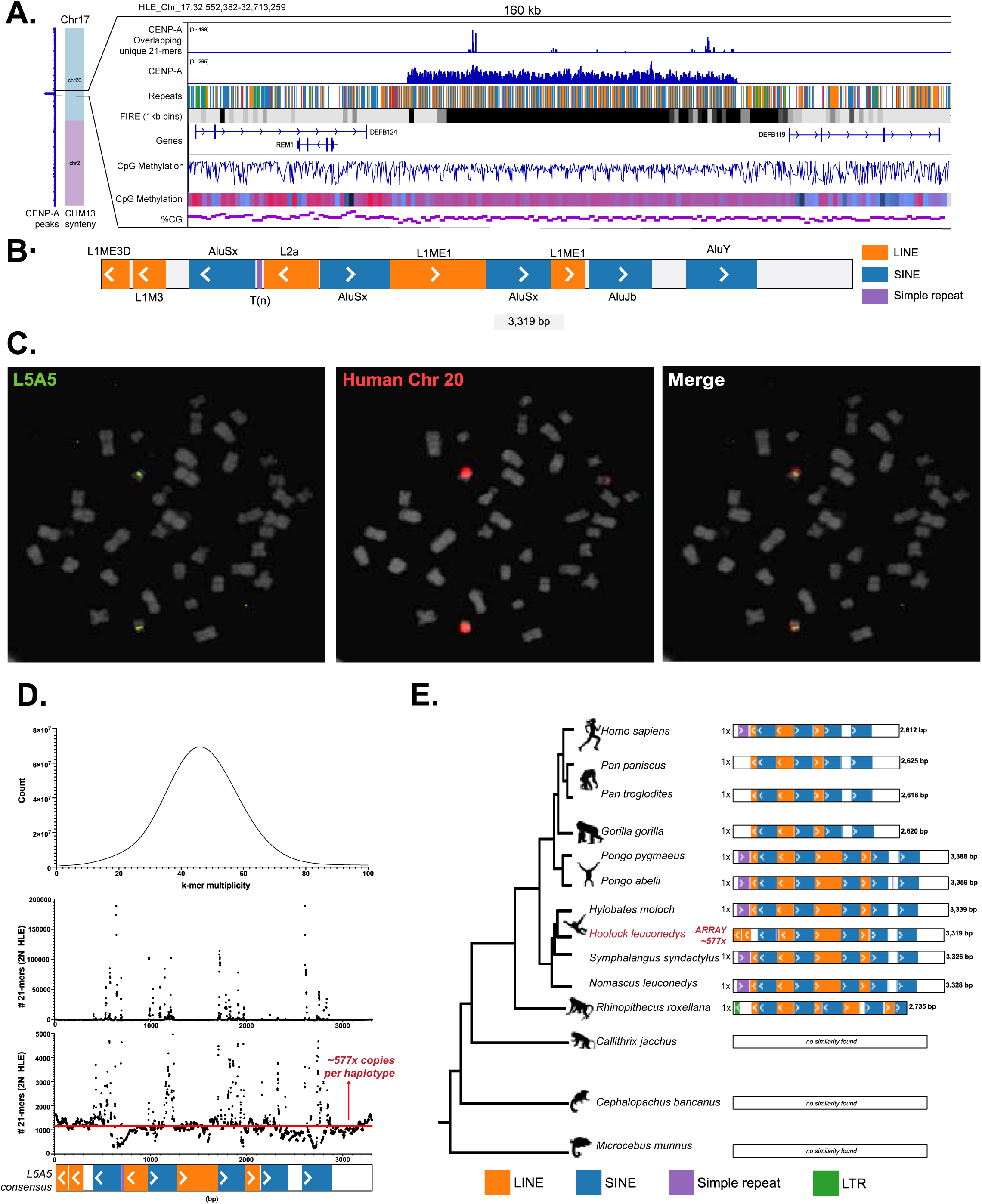
HLE Cen17 is defined by a unique composite repeat duplication not found in other apes. **(A)** To the left, HLE chromosome 17 is depicted with colors indicating synteny to T2T-CHM13. CENP-A CUT&RUN enrichment is shown vertically along the entire chromosome (blue). A zoomed panel shows CENP-A CUT&RUN mapping filtered by reads overlapping with unique 21-mers in the HLE assembly, and total unfiltered CENP-A peaks. Below, a repeat track shows the L5A5 composite repeat assembled in tandem 24 times. FIRE element density, genes, and CpG methylation, and GC percentage is shown according to the key in Figure 2. **(B)** The 3,319 bp consensus sequence of the L5A5 repeat is shown. **(C)** DNA FISH on HLE metaphase spreads using a Dig-labeled oligo specific to the L5A5 repeat shows centromeric hybridization on one chromosome pair (green). Human chromosome 20 whole chromosome paint (red) hybridizes to the same chromosome as L5A5, confirming the location of L5A5 to HLE chromosome 17, a chromosome which shares synteny to human chromosome 20^2^. **(D)** On top, distribution of 21-mer counts from PCR-free Illumina data is shown as 21-mer multiplicity (the number of times a 21-mer was found in the PCR-free Illumina reads) versus the number of 21-mers found at that multiplicity. The chart peaks at 46X, representing the estimated PCR-Free Illumina sequencing depth. Below, the L5A5 copy number is estimated. Along the x-axis, the L5A5 consensus sequence is shown, and the y-axis represents the estimated number of L5A5 repeats in the HLE diploid genome. A horizontal line represents the median of ∼1,154 copies in the HLE diploid genome (∼577 per haplotype). **(E)** A phylogeny of the L5A5 repeat across 14 primates is shown. The L5A5 repeat was found in all great apes, gibbons, and the golden snub-nosed monkey, but not in marmoset, tarsier, or lemur genomes. While the L5A5 repeat subunit structure is relatively conserved among gibbons, a SINE/*AluSx* and LINE/L1ME1 deletion shortened the consensus in great apes by ∼700 bp. HLE is the only species with an arrayed L5A5 centromeric structure; all other species have only one L5A5 copy identified.

As the L5A5 repeat has not previously been described as a centromeric repeat in other primates, we searched for copies of this repeat in 13 other primates with available high-quality genome assemblies, including representative species from each major lineage (Table S8). The L5A5 composite repeat was detected across great apes, gibbons, and the golden snub-nosed monkey (Figure 4E), but not in the marmoset, tarsier, and lemur genomes, suggesting that the L5A5 unit evolved after the split between Old and New World monkeys, likely due to active TE insertions (Figure 4E). Expectedly, L5A5 repeats in the three other gibbon genera shared the most similarity to the HLE L5A5, with one short L1M duplication at the end of the consensus differentiating them (Figure 4E). The L5A5 repeat in the golden snub-nosed monkey assembly displayed the most divergence from the *Hoolock* consensus, including additional LINE/L1ME2 and LTR/ERVL insertions and sequence deletions (Figure 4E), likely a reflection of the >25 million years divergence time between gibbons and Old World monkeys^51^. Similarly, all great apes shared a SINE/*Alu*Sx and LINE/L1ME1 sequence deletion, shortening the consensus sequence to ∼2,619 bp (Figure 4E).

While the L5A5 monomer displayed lineage-specific sequence evolution across the primate phylogeny, the most notable difference between the centromeric L5A5 repeats in HLE and the non-centromeric L5A5 loci across other primates was its copy number. While L5A5 was arrayed ∼577 times per haplotype in HLE, BLAST analysis (Figure 4E) and PCR (Figure S12, Table S9) confirmed that only one copy per haplotype was present in each of the other primate assemblies, including non-HLE gibbons. Combined with the observation that Cen17 is an evolutionary new centromere specific to *Hoolock*, these findings suggest a link between the L5A5 composite repeat amplification and the formation of the lineage-specific centromere.

### HLE breakpoints exhibit distinct genetic and epigenetic features

The availability of a high-quality, contiguous assembly for HLE, as well as the suite of ‘omics data we generated, provided a unique opportunity to investigate the genetic and epigenetic mechanisms underlying the karyotype instability in gibbons and its possible relationship with rapid centromere evolution. We therefore expanded our analysis to HLE evolutionary synteny breakpoints. We first aligned *cmHooLeu*1 against reference assemblies for human, (T2T-CHM13), *N. leucogenys* (NLE, Asia_NLE_v1), *H. moloch* (HMO, HMol_V3), and *S. syndactylus* (SSY, NHGRI_mSymSyn1-v1.1-hic.freeze_pri) to identify syntenic blocks and breakpoints using a custom python script ^52^. Overall, a total of 364 evolutionary breakpoints were identified (123, 92, 74, and 75 breakpoints comparing HLE against CHM13, NLE, HMO, and SSY, respectively, with a median size of 40.8 kb and accounting for 202 unique loci); Table S10, Table S11, Figure 5A). Among the 19 centromeres in HLE, 13 (∼68%) overlapped with predicted breakpoints (Figure 5B).

**Figure 5:**
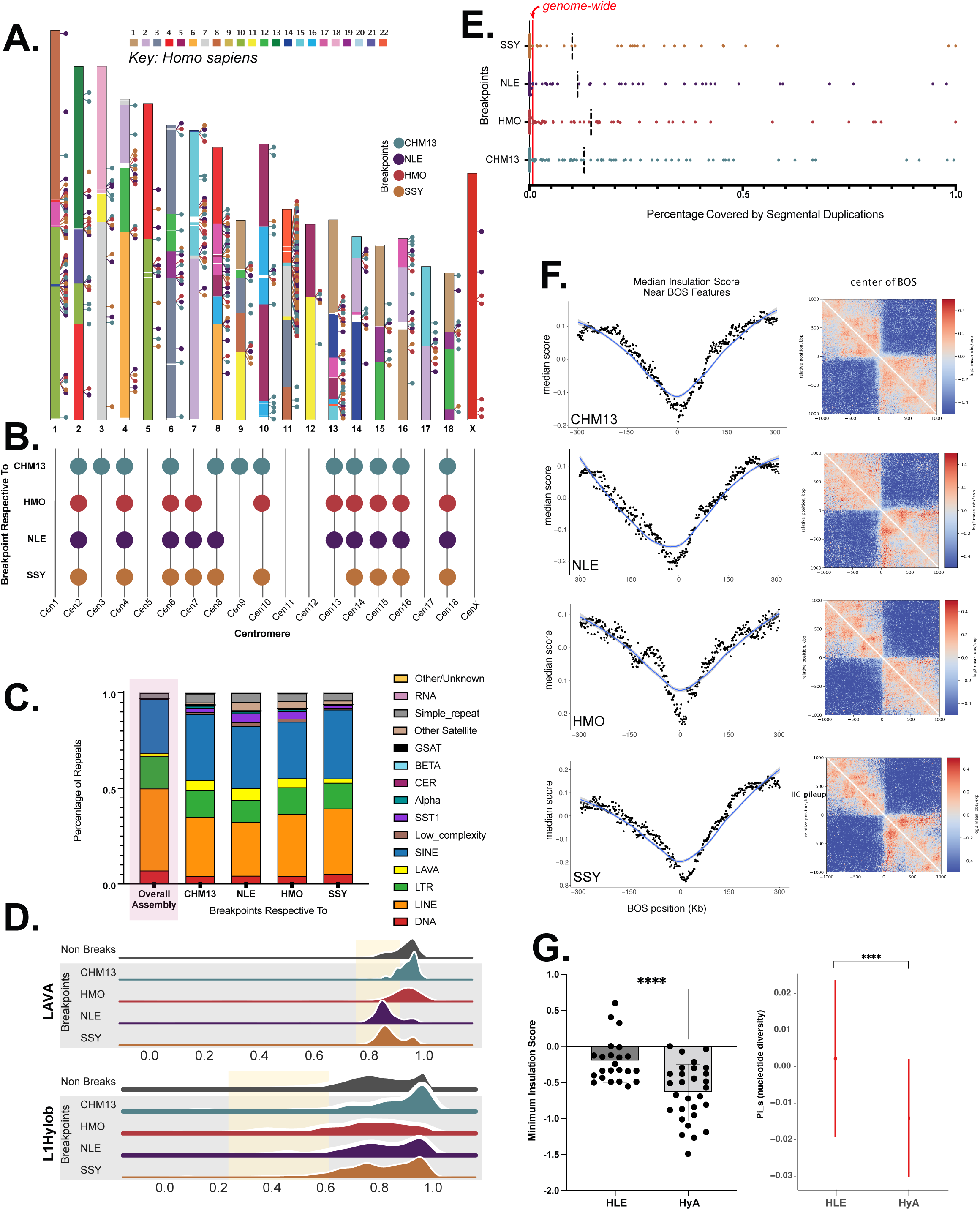
HLE breaks of synteny (BOS) exhibit distinct genetic and epigenetic features. **(A)** An ideogram of the assembled HLE chromosomes is shown, with colors corresponding to synteny between human chromosomes (T2T-CHM13) according to the key. To the right of each chromosome, circle markers indicate location of HLE BOS respective to the T2T-CHM13, NLE, HMO, and SSY genome assemblies in differing colors. **(B)** An upset plot shows BOS found at each HLE centromere. **(C)** The percentage of total repeats in the overall HLE assembly, and at BOS respective to T2T-CHM13, NLE, HMO, and SSY are shown, with each repeat class represented in a different color. SINEs, LAVAs, SST1s, and simple/low-complexity repeats are prevalent in BOS regions, while LINEs appear depleted. **(D)** Aggregated CpG methylation across LINE/L1Hylobs and LAVAs are depicted as ridgeplots, showing repeats annotated within BOS respective to CHM13, HMO, NLE, and SSY, as well as repeats outside BOS. Both L1Hylobs and LAVAs are less methylated in BOS on average (highlighted in yellow) with few exceptions. Specifically, LAVAs in HLE-NLE and HLE-SSY BOS and L1Hylobs in HLE-HMO BOS show significant shifts towards lower CpG methylation (p<0.0001). **(E)** Percentage of BOS covered by segmental duplications is shown, with one dot corresponding to each BOS and black lines indicating the average percentage of bases covered in each category (including BOS with no coverage). The vertical red line at 0.0071 indicates the coverage of segmental duplications genome-wide. **(F)** Dot plots (black) and loess smoothed curves (blue), show dips in median insulation scores at BOS. Heatmaps show reduction in the frequency of genomic interactions around BOS on a scale from low (blue) to high (red). **(G)** A dot plot of minimum insulation score (left) shows that older BOS (HyA) are more insulated (lower insulation score) than younger (HLE) BOS (p<0.0001). On the right, the marginal effect of BOS age on nucleotide diversity is plotted after controlling for other genomic features associated with nucleotide diversity. Older BOS were found to have significantly lower nucleotide diversity than younger BOS (p<0.0001). Error bars show the standard deviation.

Compared to repeat enrichment in the overall HLE assembly, breakpoints contain a higher percentage of LAVA elements (averaging 4.62% of repeats in breakpoints compared to 1.39% of the total repeats genome-wide), SST1s (averaging 3.18% of breakpoint repeats vs. 0.05% total repeats), and SINEs (averaging 33.19% vs. 28.03%) (Figure 5C, Table S12). Similarly, satellites, low-complexity repeats, and simple repeats composed a higher percentage of repeats in breakpoints than across the HLE assembly (Figure 5C, Table S12). While still prevalent in breakpoint regions, LINEs were less commonly found compared to the overall assembly, composing an average of 31.47% of repeats in breakpoints compared to 42.93% of the total repeats in the HLE assembly (Figure 5C, Table S12).

Previous work in NLE identified an enrichment of hypomethylated *Alu*s in breakpoints^53^, suggesting a correlation between epigenetic state and genome stability. CpG methylation has been shown to repress retrotransposition of TEs in mammals^54^. Therefore, the hypomethylation of *Alu*s at breakpoints may lead to higher TE activity and genome recombination in gibbons^55^. Accordingly, we assessed CpG methylation of LINEs, SINEs, and LAVAs across HLE breakpoints. Methylartist^56^ was used to plot average CpG methylation of SINEs (*Alu*Y, *Alu*Js *Alu*S), LINEs (L1Hylob, L1M, L1P), and LAVAs within and outside of breakpoints (Figure 5D, S13, Table S13). We did not detect hypomethylation of *Alu*s at breakpoints (Figure S13); in fact, on average, *Alu*s were more methylated at breakpoints than outside (Figure S13, Table S13). However, L1Hylob LINEs were less methylated at HLE-HMO breakpoints on average compared to those outside of breakpoints (p<0.0001), and LAVA elements in HLE-NLE and HLE-SSY breakpoints were less methylated than LAVAs found elsewhere in the assembly (p<0.0001) (Figure 5D, Table S13). These results suggest that hypomethylation and consequential activity of TEs may be correlated to genome instability of gibbons, yet this does not appear to be restricted to a specific repeat class.

NLE breakpoints have been found to be enriched in SDs, suggesting a link between these duplicated regions and chromosome reshuffling in gibbons^2,53,57^. We overlapped *BISER* SD predictions in the HLE assembly, filtering for SDs >1 kb in length and sharing >90% alignment identity. Of the 364 total breakpoints identified, 179 overlap with at least one SD (49.18%) (Table S14, Figure 5E). Of these, 95.5% (n=171) were covered by a higher percentage of SDs compared to the HLE assembly, for an average of 25.85% (compared to 0.71% of the total assembly SD coverage) (Table S14, Figure 5E). These observations provide additional support for a correlation between SDs in centromeres and karyotype evolution in gibbons.

Finally, several studies report an association between evolutionary breakpoint and spatial chromatin conformation, particularly boundaries of self-interacting topologically associated domains (TADs)^58–60^. Breakpoints are often found colocalized with TAD boundaries and present with reduction of chromatin interaction across the two sides of the breakpoint. Consistent with these reports, our Omni-C data showed reduction of chromatin interaction frequency at breakpoints as well as a decrease in insulation score (i.e., a measure of frequency of interactions passing through any given region of the genome) around the breakpoints (Figure 5F).Consistently, breakpoints obtained by comparing all four species against HLE were significantly closer to HLE TAD boundaries and had significantly lower median and minimum insulation score compared to random background (i.e., randomly shuffled size-matched regions in the same chromosomes; two-tailed Wilcoxon sum-rank test, p<0.05, Table S15).

In order to investigate the temporal dynamics of chromosomal rearrangements and TAD boundary establishment, we utilized previously reported FISH-based chromosome mapping^2^ to stratify breakpoints into two groups based on their evolutionary context: those shared in the ancestral karyotype state (HyA) or those found specific to the HLE lineage. We find that shared HyA breakpoints, which likely correspond to older evolutionary events, provide stronger insulation for chromatin interactions (p<0.0001) and were located closer to TAD boundaries (p<0.05) compared to younger, *Hoolock*-specific breakpoints. In addition, HyA breakpoints were also found to be associated with significantly lower estimates of nucleotide diversity than HLE breakpoints (Figure 5G, S14, S15, Table S16). Despite strong insulation and low diversity of ancestral breakpoints, no other genetic or epigenetic features were found to be significantly different among these breakpoints, including breakpoint size, segmental duplication coverage, CpG methylation, replication timing, chromatin accessibility, predicted CTCF binding sites, gene content or repeat composition (with the exception of LAVA, which was found to be slightly more present in older breakpoints) (Figure S16). Combined, these features suggest that older breakpoints are more constrained regardless of DNA content or other epigenetic features and thus represent stronger TAD boundaries.

## Discussion

While most DNA sequences involved in essential cellular functions are highly conserved across species, centromeric DNA and associated proteins evolve rapidly, indicating that centromeres are specified and maintained through epigenetic mechanisms^61^; yet, the factors driving centromere localization and function have been difficult to elucidate using short-read based methods. However, advancements in centromere assembly have now afforded the opportunity to map key epigenetic features that distinguish functional centromeres. In this study, we used long-read sequencing to assemble the eastern hoolock gibbon (HLE) genome and generate six gapless gibbon centromeres, enabling the first –to our knowledge– genome-scale analysis of gibbon centromere content and organization at the level of single chromosomes. We report that HLE centromeres vary in their TE and satellite composition across chromosomes, differing from the canonical alpha satellite organization of other ape centromeres^8^. Despite this difference, functional HLE centromeres retain a CpG methylation dip concomitant with CENP-A nucleosome enrichment (centromere dip region, CDR), indicating that this epigenetic feature is conserved across ape lineages and is independent of alpha satellites. Additionally, we find that dichotomous compacted and accessible chromatin (“dichromatin”) is conserved among both human alpha-satellite arrays and the TE-rich centromeres. Thus, the conserved epigenetic structure of primate centromeres includes centromere-specific histones (CENP-A), dichotomous chromatin, and a CDR independent of sequence type (satellite vs TE vs composite repeat) or organization (complex vs HOR vs array).

Despite conservation of epigenetic structure, we find several features that differentiate gibbon centromeres from their human counterparts. We identified an overlap between an evolutionary breakpoint in the *PLSCR* protein-coding gene cluster and CENP-A enrichment in HLE Cen9, a significant dissimilarity between the gene-poor heterochromatic regions underlying CENP-A bound regions in humans and many other species. While *PLSCR2* is lowly expressed in humans^62^, in gibbons, lack of detected expression paired with its recombined, potentially inactive structure due to chromosomal rearrangement and localization at a HLE centromere suggests *PLSCR2* loss of function. In addition, we identified the presence of *CLUHP* pseudogenes directly overlapping with CENP-A enrichment as well as other genes associated with enrichment near centromere domains (*CDK5RAP3*, *EME1*, *DEFB119*, *DEFB4A*). Notably, these genes have paralogs and pseudogenes sharing high sequence identity between multiple chromosomes, perhaps contributing to rearrangements via non-allelic homologous recombination events.

DNA replication within the cell during S phase is a highly tuned, dynamically regulated temporal process^63,64^. Replication timing domains, functional units of replication origins that initiate synchronously, provide a characteristic and coordinated temporal organization for DNA replication dependent on cell and developmental status^63,64^. It has been reported that centromeres have variable replication timing across eukaryotes, with centromeres replicating in early S phase across yeast species^65–68^, mid S phase in maize^69^, and mid to late S phase in humans^13,70–73^; yet, within a species, the replication timing of centromeres is conserved among chromosomes. Replication stress has been shown to be caused by several factors, including chromatin compaction, collisions between transcriptional and replicative machinery, repeats, secondary structures, and histone modifications (reviewed in ^74^). In fact, most active genes and SINE/Alus are early replicating, while LINEs and centromeres have been reported to be late replicating^74,75^. **S**INE-R, **V**NTR, ***A****lu*-like (SVA) elements, which share the VNTR and *Alu*-like components with LAVA, have been shown to cause replication fork stalling leading to later replication^76^. Such stress has been linked extensively to genome instability in mice and humans^47^. Here, we found discrepancy in replication timing of HLE centromeres compared to humans, which take place in mid-late S phase^71^. Instead, HLE chromosomes contain centromeres with both early and late replicating DNA. We postulate that a source of such asynchrony lies within the highly variable repeat composition of the HLE centromere. Moreover, the three HLE centromeres with early S phase replication (Cen8, Cen9, and Cen11) all have genes overlapping or directly upstream of CENP-A enrichment regions, significant upstream hypomethylation concomitant with increased chromatin accessibility, and pericentromeric LAVA expansions comprised of TEs with likely opposing replication timing expectations. Hence, such conflicting features may cause replication stress and variation in replication timing, which may in turn lead to genome instability. However, whether replication stress is a cause or consequence of the chromosome instability observed in HLE (and possibly in all gibbons) is still yet to be determined.

In addition to replicative stress at the centromeres, our analysis of evolutionary breakpoints in the HLE genome suggests an additional mechanism that may contribute to genome instability. CpG methylation is an important regulator of mobile element activity, suppressing TE expression and subsequent propagation in the genome, which can drive chromosomal rearrangements^54,77^. Previous studies in *Nomascus leucogenys* (NLE) showed that *Alu* elements at breakpoints were less methylated than those outside of breakpoints, suggesting these epigenetic differences contribute to TE activity leading to repeat-mediated chromosomal rearrangements^53^. While we did not detect similar patterns of *Alu* hypomethylation at breakpoints in HLE, we find hypomethylation of other TEs, particularly /L1Hylobs and LAVAs, at breakpoint regions. Given the high rates of LAVA proliferation in HLE compared to other gibbon genera, it is clear that TEs have undergone unique evolutionary histories since the radiation of gibbons from a common ancestor. Therefore, it is plausible that similar to *Alu*s in NLE, hypomethylation of L1Hylob and LAVA elements contributed to karyotype diversity in the *Hoolock* genus.

Finally, we extended our analysis of breakpoints into topologically associated domains (TADs), chromatin domains that serve as fundamental units of three dimensional genome organization and are hypothesized to serve as gene regulatory units by controlling long-range interactions^78^. The conservation of such domains across species has been associated with the conservation of syntenic regions across evolution, and studies have identified that chromosomal breakpoints are more common at TAD boundaries than inside of TADs^58,60,79,80^. These observations have reinforced the hypothesis that TADs are evolutionarily constrained, and chromosomal rearrangements disrupting TADs are negatively selected. Using Omni-C data, we found a reduction of chromatin interaction frequency and insulation scores at breakpoints, concordant with previous findings suggesting breakpoints coincide with TAD boundaries^52,60^.

Additionally, we find that ancestral breakpoints among gibbons generate stronger interaction insulation and have lower nucleotide diversity than younger breakpoints despite no other significant differences in genetic and epigenetic features (with the exception of LAVA elements). Collectively, these data suggest that there is a relationship between genome rearrangements and maintenance of genome topology. Combined with the observations in HLE centromeres described above, repeat-mediated chromosomal rearrangements paired with a “perfect storm” of centromeric replicative stress emerges as a potential driver for karyotypic variation seen in gibbons. We therefore propose that an array of interconnected epigenetic and genetic features, rather than just one isolated element, contributes to the genome remodeling observed in gibbons. All features identified within and around HLE centromeres, including hypomethylation, chromatin compaction, transposable elements, SDs, and satellite arrays, have been linked to genomic instability in other model systems and other gibbon species. We speculate that these various features all impose unique replicative stress on the HLE genome, as the finely tuned replication timing program has to balance the presence of centromeric coding genes and associated transcriptional machinery, extreme variation in methylation and chromatin compaction, as well as large arrays of tandemly repeated SST1 and LAVA segmental duplications. Continued efforts to produce high-quality genome resources from gibbons promise to unravel the mechanisms dictating their unique chromosome evolution and provide much needed genomic information for conservation management efforts for these endangered apes.

## Data Availability

Assembled genome and sequencing data are available under BioProject ID PRJNA1153068 and BioSample ID SAMN43386187.

## Acknowledgements

R.J.O and G.A.H. were supported by NIH R01GM123312 and NSF 2217100. L.C. is supported by grants from the National Institutes of Health, R01 HG010333 and P51 OD011092 (to the Oregon National Primate Research Center). D.M.G. is supported by National Institutes of Health R01 HG007352. The authors would like to acknowledge the Center for Genome Innovation for sequencing resources and the UConn Computational Biology Core for computational resources. The authors would also like to acknowledge Gabriel Mirasol for the gibbon illustrations depicted in Figure 1. Finally, the authors would like to acknowledge the Gibbon Conservation Center in Santa Clarita, California, for their sample support and continued efforts in gibbon conservation.

## Author Contributions

Conceptualization: GAH, LC, RJO; Data curation: GAH, RJO; Formal analysis – genome assembly: GAH, RJO; Formal analysis – repeat and segmental duplication annotation: GAH, JMS, RJO; Formal analysis – fluorescence in situ hybridization: GAH, RJO; Formal analysis – alpha satellite annotation: IA, FR; Formal analysis – substitution rates: NA; Formal analysis – gene predictions: EF, WT; Formal analysis – breakpoint and TAD analysis: GAH, MO, JW, LC; Formal analysis – FIRE: GAH, SN, DD, AS; Sequencing Data generation – genome sequencing (ONT): NP, GAH, RJO; Sequencing Data generation – genome sequencing (PacBio/Fiber-seq): GAH, YM, AS, RJO; Sequencing Data generation – genome sequencing (Illumina PCR Free): NP, GAH, RJO; Sequencing Data generation – Omni-C: CM, NP, RJO; Sequencing Data generation – PRO-seq: SJH, RD, DS, LC; Sequencing Data generation – Repli-seq: GAH, TS, DMG, RJO; Funding acquisition: LC, RJO; Sample acquisition: LC, RJO; Writing – primary: GAH, RJO; Writing – review & editing: GAH, MO, SJH, EF, RJO.

## Methods

### Oxford Nanopore Technologies (ONT) Sequencing

Cells were collected from a transformed *Hoolock leuconedys* lymphoblastoid cell line (LCL). High molecular weight (HMW) genomic DNA was extracted following a modified protocol^83^. In brief, cells were lysed in 300ul lysis buffer (400 mM Tris pH 8.0; 60 mM EDTA pH 8.0; 150 mM NaCl; 1% SDS with 15 uL Puregene Proteinase K (Qiagen)) at 55°C for 2 hours with inversion every 30 minutes. Post incubation, an additional 185 uL of lysis buffer was added and incubated overnight at 50°C. The lysate was treated with 500ug/ml of RNaseA, incubated at 37°C for 20 minutes. The DNA was extracted using an equal volume of phenol:chloroform: isoamyl alcohol (25:24:1), followed by two rounds of chloroform: isoamyl alcohol (24:1). The aqueous layer containing the DNA was collected and the DNA precipitated with ethanol, DNA was spooled and washed with 70% ethanol. The DNA was resuspended in nuclease free water. Library preparation was performed using the Ligation Sequencing Kit (LSK109). HMW DNA was sequenced on the PromethION platform from Oxford Nanopore Technologies using a PromethION R9.4 FLOPRO002 flow cell and basecalled using Guppy (v2.2.3)^84^. Across five flow cells, a total of 89 Gb passed quality filtering (∼31x coverage). Before assembly, obtained reads were recalled using Guppy (v5.0.16)^84^ to improve basecalls.

To isolate ultra-high molecular weight (UHMW) DNA, the Circulomics Nanobind UHMW DNA Extraction for cultured cells (EXT-CLU-001) protocol was followed according to manufacturer’s instructions with the Circulomics Nanobind CBB Big DNA Kit (NB-900-001-01). Library preparation was performed using the Circulomics Nanobind Library Prep protocol for ultra long sequencing (LBP-ULN-001) using the Circulomics Nanobind UL Library Prep Kit (NB-900-601-01) and the Oxford Nanopore Technologies Ultra-Long DNA Sequencing Kit (SQK-ULK001). UHMW DNA was sequenced on the PromethION platform using a R9.4 FLOPRO002 flow cell and basecalled using Guppy (v5.0.16)^84^. A total of 45 Gb passed quality filtering (∼16x coverage).

### Illumina PCR-free Sequencing

In order to generate highly accurate PCR-free Illumina sequencing reads used for polishing and QV score estimation, the Illumina DNA PCR-Free Library Prep Tagmentation protocol was followed for library preparation according to the developer’s instructions. The library was sequenced on the Illumina NovaSeq 6000 platform to a depth of ∼50X. Reads were QCed using FastQC (v0.11.7)^85^ and trimmed using cutadapt (v3.5)^86^ prior to genome polishing using a quality cutoff score of 20 (-q 20) and a minimum read length of 50 bp (-m 50).

### Dovetail Omni-C Sequencing

To generate Dovetail™ Omni-C™ reads, roughly 1.5 million cells were collected from the previously described *Hoolock leuconedys* LCLs and processed according to the Dovetail™ Omni-C™ Proximity Ligation Assay protocol for mammalian samples (v1.4). Lysate quantification was performed using the Qubit® dsDNA HS kit for the Qubit® Fluorometer and the D5000 HS kit for the Agilent TapeStation 2200. 150 bp paired end sequencing was performed on the Illumina NextSeq 550 V2 platform to a depth of ∼274M reads.

### Genome Assembly and QC

Full assembly code can be found on Zenodo^87^. Flye (v2.9)^88^ was used to assemble the raw Oxford Nanopore reads using an estimated genome size of 2.9 Gb, the size of the previously assembled *Nomascus leucogenys* genome. Medaka (v1.4.3)^89^ was used for long read polishing using default settings and the r941_prom_sup_g507 model. Publicly available Illumina WGS sequencing reads for the same individual (SAMN12702557) were used to polish the assembly by mapping the reads with Burrow’s Wheeler Aligner (v0.7.17)^90^ using the bwa mem algorithm and processed using Samtools (v1.7)^91^. Short read error correction was performed using Pilon (v1.22)^92^ with default parameters. Haplotype redundancies and assembly artefacts based on read coverage were removed using minimap2 (v2.15)^93^ and PURGEhaplotigs (v1.0)^94^. The reformat.sh module of BBMap^95^ was used to impose a 3kb limit on the genome.

Omni-C™ sequencing quality control was performed using FastQC (v0.11.7)^85^ and was analyzed according to the Dovetail™ documentation using the Dovetail’s pre-built environment. Briefly, Burrow’s Wheeler Aligner (v0.7.17-r1188)^90^ and Samtools (v1.9)^91^ was used to align and process the Omni-C™ reads to the assembly using the -5SP and -T0 flags to accommodate independently mapping mate pairs. Pairtools (v0.3.0)^96^ was used to identify valid ligation events (pairtools parse; –min-map 40, --walks-policy 5unique, and –max-inter-align-gap 30 flags), sort the file (pairtools sort) remove PCR duplicates (pairtools dedup; -mark-dups and -output-stats flags) and split the file (pairtools split; -output-pairs and -output-sam flags). Dovetail’s get_qc.py script was used to retrieve key library statistics and Preseq (v0.1.24)^97^ was used to estimate library complexity.

Juicer (v1.6)^98^ and 3D-DNA (v180922)^99^ were used to scaffold the assembly using the protocols outlined by developers. Juicebox with Assembly Tools (v1.11.08)^100^ was used for manual review of the produced scaffolds. LASTZ (v1.04.15)^101^ and UCSC GenomeBrowser Kent Tools (v369)^102^ were used to align the assembly to the CHM13 v1.1 genome using a custom pipeline (https://github.com/carbonelab/lastz-pipeline). Resulting alignments were validated according to predicted syntenic regions and large-scale chromosome misassemblies and misorientations were manually corrected using Emboss (v6.6.0)^103^.

To reduce any misassemblies associated with manual curation and scaffolding, pre-scaffolded contigs (the “query”) were aligned to the curated assembly (the “reference”) and scaffolded using RagTag (v2.1.0)^104^. The resulting scaffolded assembly (built from “query” contigs) was gap filled using TGS-GapCloser (v1.0.1)^105^. The gap filled, final assembly was polished with Illumina reads using Pilon (v.1.22)^92^ with default parameters.

Quality metrics of the assembly were analyzed using QUAST (v5.0.2)^106^. BUSCO (Benchmarking Universal Single-Copy Orthologs) (v5.0.0; MetaEuk v4.0)^107^ was used to assess assembly gene completeness using the lineage dataset primates_odb10. To assess *k*-mer completeness, Meryl (v1.3)^18^ was used to count 21-mers found in the Illumina PCR-free library, and Merqury (v1.3)^18^ was used to determine the QV score and estimate completeness of identified 21-mers within the assembly. GenomeScope2.0^108^ was used to estimate the HLE genome size.

### Repeat, CpG Methylation, Gene, and Segmental Duplication Annotations

Repeats in the genome were annotated with RepeatMasker (v4.1.2-p1)^109^ using the Crossmatch search engine (v1.090518)^110^ and a combined gibbon (*Hylobates* sp.) Dfam (v3.6) and Repbase (20181026) repeat library with the “-lib gibbon” flag. Repeats identified in the CHM13 genome not yet included in the gibbon Dfam repeat lineage^21^ were annotated using a custom repeat library against the draft masked genome. The two repeat annotations were merged using the RMComp.pl pipeline^21^. The RepeatMasker summary script buildSummary.pl was used to summarize the percent count and percent base pairs of each repeat. CpG methylation was called using Bonito (v2.1.2)^111^ in raw ONT reads. Raw reads were converted to Fastq format using Samtools (v1.9)^91^, and mapped to the HLE assembly using Winnowmap (v2.03)^112^. Modbam2bed (v0.9.5)^113^ was used to generate viewable aggregated CpG methylation tracks. Genes were predicted and annotated using FLAG^114^.

Segmental duplications in the HLE assembly were predicted using *BISER* (v1.4)^40^ using default parameters. Next, segmental duplications were filtered using awk to include only predictions that both 1) were >1 kb in length and 2) shared >90% ungapped sequence identity. Segmental duplication predictions were overlapped with *RepeatMasker* LAVA and SST1 annotations using Bedtools (v2.29.0) intersect^115^. Segmental duplications were visualized using Circos (v0.69-9)^116^.

### CENP-A CUT&RUN Preparation, Sequencing, Analysis, and Validation

The CUT&RUN Assay Kit (#86652) from Cell Signaling Technology® was used to assess CENPA protein-DNA interactions following manufacturer’s instructions. 250,000 cells per condition were pelleted and washed in 1X wash buffer (10X wash buffer [#31415], 100X spermidine [#27287], 200X protease inhibitor cocktail [#7012]). Cells were bound to Concanavalin A beads for 5 minutes at room temperature, then resuspended in 1X binding buffer (100X spermidine, 200X protease inhibitor cocktail, 40X digitonin solution [#16359], and antibody binding buffer [#15338]). To assess CENPA-DNA interactions, the CENP-A monoclonal antibody (Enzo, ADI-KAM-CC006-E) was used at a dilution of 1:50. Tri-methyl-histone H3 (Lys4) (Cell Signaling Technology, C42D8) rabbit monoclonal antibody (#9751) at a dilution of 1:50 was used as a positive control; rabbit (DA1E) monoclonal antibody IgG XP® isotype control (Cell Signaling Technology, #66362) at a dilution of 1:10 was used as a negative control. The antibodies were bound at 4°C for 2 hours, then the beads were washed in digitonin buffer (10X wash buffer, 100X spermidine, 200X protease inhibitor cocktail, and 40X digitonin solution) on a magnetic rack. The beads were resuspended in digitonin buffer and pAG-MNase enzyme (#40366) then incubated at 4°C for 1 hour. Following, the beads were washed in digitonin buffer on a magnetic rack, then resuspended in digitonin buffer and calcium chloride and incubated at 4°C for 30 minutes. Digestion was stopped with 1X stop buffer {4X stop buffer [#48105], digitonin solution, and 200X RNase A [#7013]). For normalization, 10 pg/uL spike-in DNA [#40366] was added at a 1:100 dilution. The samples were incubated at 37°C for 10 minutes, then supernatants were transferred to a new microfuge tube. Samples were incubated at 65°C for 2 hours before proceeding to DNA purification. Input chromatin samples were sheared to ∼100-700 bases using a Covaris S2 sonicator prior to purification.

DNA purification was performed using the Cell Signaling® DNA purification with spin columns kit (#14209). DNA concentration was assessed using the Qubit® dsDNA HS kit for the Qubit® Fluorometer and the High Sensitivity D1000 kit for the Agilent TapeStation 2200. CENP-A and Input libraries were prepared using the NEBNext Ultra II DNA Library Prep Kit for Illumina (#E7645S) and sequenced using the Illumina NovaSeq 6000 150 bp paired end settings to a depth of ∼15M reads.

Quality of the CENP-A enriched chromatin and input sequencing was analyzed using FastQC (v0.11.7)^85^. Reads were trimmed using Cutadapt (v3.5)^86^ using a quality cutoff score of 20 (-q 20) and a minimum read length of 50 bp (-m 50). Trimmed reads were aligned to the HLE assembly using Burrows-Wheeler Aligner (BWA) (v0.7.5a-r405)^90^ using a minimum seed length of 50 bp (-k 50) and skipping seeds with more than 1 million occurrences (-c 1000000). Alignments were filtered to remove multi-mappers (-F 2308) using Samtools (v1.9)^91^ and converted to bed format using Bedtools (v2.29.0)^115^. Because of the repetitive nature of centromeres, a marker-assisted filtering approach was implemented to retain only aligned reads that overlapped with a unique *k*-mer in the assembly. Meryl (v1.3)^18^ was used to generate a database of 21-mers from the HLE assembly (meryl k=21 count), filter the resulting assembly for unique *k*-mers (meryl equal-to1) and convert the database to a bed file (meryl-lookup -bed). The overlapSelect module of GenomeBrowser tools (v20180626)^102^ was used to intersect CENP-A and input alignments with unique 21-mers. The resulting alignments, now containing only reads that overlap a unique 21-mer in the assembly, were converted to bedgraphs using Bedtools (v2.29.0) ^115^ for viewing in IGV. To validate assembled centromeres, read coverage over regions with CENP-A enrichment were analyzed using NucFreq (v0.1)^50^. Of the 19 assembled chromosomes, 13 had anomalies in read coverage; therefore, the six well assembled centromere regions were targeted for further analysis.

### Assessment of Repeat Content, Methylation, and Genes Across Active Centromeres

In order to assess the repetitive content across each of the six uncollapsed HLE centromeres, the *RepeatMasker* annotations were converted to bed format and intersected with regions of CENP-A enrichment using Bedtools (v2.29.0) intersect^115^. Overall repeat composition across CENP-A regions was summarized using the buildSummary.pl utility script within the *RepeatMasker* package^109^. Repeat tracks were visualized using Integrated Genomics Viewer (IGV). Sequence similarity was assessed using StainedGlass (v0.5)^117^. CpG methylation tracks were generated using Modbam2bed (v0.9.5)^113^. Detailed visualization of CpG methylation across centromeres was performed using the methylartist locus command available with Methylartist (v1.2.7)^56^. Gene predictions were overlapped with CENP-A regions using Bedtools intersect (v2.29.0)^115^ and gene ontology was examined using the GO database^118,119^.

To validate the gene predictions that overlap with CENP-A enrichment on HLE chromosome 8 and chromosome 9, these regions were aligned to homologous genes from human (T2T CHM13v2.0/hs1), Rhesus macaque (Mmul_10/rheMac10), marmoset (Callithrix_jacchus_cj1700_1.1/calJac4), and NLE (GGSC Nleu3.0/nomLeu3) using Clone Manager Professional 9 software. For the *CLUHP* pseudogenes predicted at HLE Cen8, exons were manually annotated based on all 11 *CLUHP* genes annotated in NCBI for human and additional human *CLUHP* genes identified using the BLAT tool in the UCSC browser. For the *PLSCR1*, *PLSCR2*, and *PLSCR2*-like genes predicted on HLE chromosome 6 and at HLE Cen9, exons were manually annotated based on homologous exons identified in human, macaque, marmoset, and NLE.

### Single-molecule chromatin fiber sequencing and processing

To perform Fiber-seq, cells were collected and processed according to the methods in ^14^. Reads were processed with jasmine then passed into the FIRE pipeline^120^ to call inferred regulatory elements (FIREs) and binned to 1kb for visualization.

### Alignment of RNA-sequencing to HLE Assembly

Publicly available RNA-sequencing was mapped to the HLE assembly using Hisat2 (v2.2.1)^121^. Reads were processed to bam format using Samtools (v1.9)^91^ and converted to bedgraph format for visualization using Bedtools (v2.29.0) bamtobed and Bedtools (v2.29.0) genomecov^115^.

### Precision Run-On sequencing (PRO-seq) and analyses

At the time of harvest, 8×10^6^ cells were collected and pelleted in a swinging bucket centrifuge at 800 x g for 5 min at 4°C. Media was aspirated without disturbing the cell pellet followed by washing with 1x PBS, pipetting gently to break up the cell pellet. After another spin, the PBS was aspirated and 4 ml of cold Chromatin Lysis Buffer (20mM HEPES pH 7.5, 300 mM NaCl, 0.2 mM EDTA, 1M Urea, and 1% NP-40; a final concentration of 1mM DTT, 0.01% (w/v) Dextran Sulfate MW 500,000, and RNase Inhibitor were added prior to use) was added directly to the cell pellet, mixing by pipetting. Samples were incubated on ice for 5 min before being transferred to a 5mL tube and spun in a swinging bucket rotor at 2,500 x g for 8 min at 4 °C. Chromatin Lysis Buffer was aspirated and the chromatin pellet was washed twice with 4 mL of cold Chromatin Wash Buffer, (150mM KCl, 10mM Tris-Cl pH 8, 10% Glycerol, 250mM Sucrose, 500 mM Betaine; 0.5mM DTT and RNase Inhibitor were added prior to use.) mixing with a gentle vortex prior to being spun at 7,500 x g for 8 min at 4 °C. Chromatin Wash Buffer was then aspirated and 0.5 mL of Buffer F (50 mM Tris-Cl pH 8, 40% Glycerol, 1 mM EDTA, 5 mM MgCl; 1mM DTT and RNase Inhibitor were added prior to use) was added and the chromatin pellet was transferred to a 1.5 mL tube before being spun at 7,500 x g for 8 min at 4°C. After pelleting, 0.4 mL of Buffer F was removed and the chromatin pellet was snap-frozen prior to being stored at −80°C.

PRO-seq libraries were generated using biological replicates of chromatin by following the protocol described in Mahat et al^20^. and Judd et al.^122^, with the following modifications. Four biotin run-on reactions were carried out in a final volume of 200 uL and 25,000 permeabilized Drosophila S2 nuclei were added as a spike-in control during the reaction. During the run-on reaction, samples were vigorously pipetted for 60 seconds after adding the run-on master mix, then incubated at 37°C for 10 min. After the run-on, RNA was extracted using Norgen RNA purification columns (Cat #37500) as per the manufacturer’s protocol. Following purification, RNA was base-hydrolyzed for 20 min on ice prior to enrichment of biotinylated nascent RNA using 25 uL MyOne Streptavidin C1 DynaBeads (Invitrogen, Cat #65001). 3’ RNA adapter ligation was carried out off-bead using a 15 pmol adapter, followed by 5’ decapping, 5’ hydroxyl repair, and 5’ RNA adapter ligation performed on beads. Upon completion of reverse transcription, libraries were pre-amplified for 5 cycles using cycling parameters from ^20^. Test amplifications using serial dilutions of the pre-amplified libraries were then performed to determine the ideal number of cycles for full-scale amplification, with 11 cycles chosen for both samples. Fully amplified libraries were purified using NEB Monarch PCR & DNA cleanup kits (Cat # T1030L, quantified by Qubit, pooled in an equimolar fashion, and submitted for paired-end sequencing on an Illumina NextSeq 500 at the Center for Genome Innovation (UConn, Storrs, CT). Sequencing reads were partitioned such that read 1 was 44 base pairs (bp) in length, while read 2 was 40 bp. A total of three sequencing runs were performed to reach the desired sequencing depth of 70 million and 81 million paired end reads for replicate 1 and replicate 2, respectively.

Raw fastq files were first trimmed for quality (--nextseq-trim=20) and adapter sequences and then trimmed to a length of 44bp (--length) before discarding any remaining reads <15bp (-m) using cutadapt (v3.5)^86^. For this study, a three-way quantification of transcription was applied to capture the range of possible transcriptional activity across the genome, mimicking the approach laid out by ^21^.

For alignment with Bowtie2 (v2.5.0)^123^, paired-end reads were mapped to a combined HLE -Drosophila assembly (NCBI RefSeq assembly: GCF_000001215.4) using default “best match” parameters along with parameters to prevent the reporting of discordant mate (--no-discordant) and individual mate (--no-mixed) alignments, hence retaining only confident alignments. Sam files containing reads mapped to HLE were processed into a bed file for both plus and minus strands using Bedtools (v2.29.0)^115^. This bed file was subsequently used for either: 1) counting read abundance across repeats with BEDtools, or 2) generation of a single-nucleotide 3’ end only BigWig file^102,115^ indicating RNA polymerase occupancy. The latter was used for visualization in the UCSC genome browser and for heatmap generation of genic transcriptional activity genome-wide with deepTools^124^.

For alignment with Bowtie (v1.3.1)^123^, read 1 was reverse complemented using seqkit (v2.2.0)^125,126^ prior to being mapped to a combined HLE - Drosophila assembly (NCBI RefSeq assembly: GCF_000001215.4) using -k 100 parameters, reporting up to 100 valid alignments per read, and zero mismatches (i.e. a perfect alignment). Specifying zero mismatches for Bowtie2 (above) was not required as this is already the default behavior. Following mapping, sam files containing reads mapping to HLE were processed into a bed file for both plus and minus strands using Bedtools (v2.29.0)^115^. This bed file was then subsequently used for either: 1) unique 21-mer filtering, 2) counting read abundance across repeats with BEDtools, or 3) BigWig file generation of the 3’ end (Bedtools (v2.29.0)^115^, GenomeBrowser/20180626) for visualization in the UCSC genome browser.

Single-copy 21-mers were generated from the HLE assembly using Meryl (v1.3)^18^. Bed files of the Bowtie -k 100 mapped paired-end reads were used to filter through Meryl single copy 21-mers using overlapSelect with the option ‘-overlapBases=XXbp’ (wherein, XX represents the length of the single copy k-mers (21-mer)^102^. This location-based filtering method requires that a minimum of the entire length of the k-mer (21bp in this study) should overlap with a given read in order to be retained.

To assess the similarity and variation between the two PRO-seq replicates, a principal component analysis (PCA) plot was generated from Bowtie2 (default, “best match”) position sorted bams using deepTools (v3.5.0)^124^. The spearman correlation was visualized based on the output of multiBamSummary. After confirming the strong correlation between the two replicates, they were merged together for all other plots and visualizations. Heatmaps representing PRO-seq transcriptional profiles of genes were generated with deepTools computeMatrix and plotHeatmap^124^. Specific plotting parameters include: --averageTypeBins max and -- averageTypeSummaryPlot mean, and --zMax 6. PRO-seq read counts across each repeat class was obtained with BEDtools coverage -counts, requiring at least half the read pair to overlap a given repeat in order to be reported^115^.

### Assessment of replication timing using E/L Repli-seq

To assess replication timing in HLE, 5 million HLE suspension cells were labeled with BrdU (Sigma-Aldrich #B5002) to a final concentration of 100 μM. Cells were incubated for 2 hours at 37°C to allow for BrdU incorporation, then cells were pelleted at 1000 rpm for 5 minutes. The supernatant was discarded, and 2.5 mL of cold PBS/1% FBS was added. Cells were gently mixed and transferred to a round bottom tube. 7.5 mL of cold 100% ethanol was added with gentle shaking and stored at −20°C before cell sorting.

Following BrdU labeling, cells were processed according to the E/L repli-seq protocol by ^49^ but with some modifications. In short, nuclei were obtained by pepsin treatment as described in the supplementary methods in ^49^ then stained with propidium iodide (Sigma-Aldrich, #P4170) to assess cell cycle phase, and sorted by fluorescence-activated cell sorting (FACS) into early-S and late-S fractions according to the DNA content. DNA was prepared from each pool independently using the Zymo Quick-DNA Microprep kit (Zymo, #D3021), then purified DNA was fragmented and adaptor-ligated with NEBNext Ultra II FS kit (NEB #E7805) according to the manufacturer’s instructions. Finally, adaptor-ligated DNA fragments were immunoprecipitated with an anti-BrdU antibody (BD, #555627) and anti-mouse secondary antibody (Sigma-Aldrich #M7023) to capture BrdU-labeled DNA. DNA was indexed with NEB Multiplex Oligos for Illumina (NEB, #E7600S), and purified using AMPure XP beads (Beckman Coulter #A63880). Libraries were sequenced using the Illumina NovaSeq 6000 platform to generate 150 bp paired-end reads.

Early (E) and late (L) S phase Repli-seq data was processed according to the protocols in ^49^. First, Bedtools (v2.29.0) makewindows^115^ was used to make 5 kb windows corresponding to the HLE assembly. Bowtie2 (v2.5.0)^123^ was used with the –no-mixed –no-discordant –reorder -X 1000 flags, then alignments were processed, sorted, and filtered using Samtools (v1.9)^91^ to retain alignments with a MAPQ score greater than 20. Samtools (v1.9) rmdup^91^ was used to remove duplicate reads, then Bedtools (v2.29.0) bamToBed and Bedtools (v2.29.0) intersect^115^ were used to intersect the alignments with the generated 5 kb windows. The coverage was assessed using a custom script in ^49^, and the base 2 log ratio of early versus late S-phase samples were calculated over the 5 kb genomic windows. Finally, samples were post-processed using quantile normalization and Loess smoothing using the preprocessCore package. Final bedgraphs were visualized in IGV.

### Alpha satellite subfamily (SF) annotation

Alpha satellite SF were annotated according to the methods in ^27^. To align HLE alpha satellites to SSY alpha satellites, a dot matrix was built using Gepard^82^ with word length 50 and window size 0. Alpha satellites were aligned to genomes previously describe ^27,127^

### L5A5 oligo probe design and fluorescence *in situ* hybridization

To create a consensus sequence for the L5A5 repeat, each of the 24 assembled L5A5 units were manually obtained from IGV and imported into the Geneious software (build 2021-07-19 12:20)^128^. The 24 sequences were aligned to one another using a Geneious global alignment with free end gaps using a cost matrix of 65% similarity, a gap open penalty of 12, a gap extension penalty of 3, and 2 refinement iterations. From the resulting alignment, a consensus sequence was generated using a 0% majority threshold (the most common base at each position). The resulting 3,319 bp consensus sequence was annotated using RepeatMasker (v4.1.2-p1)^109^. BLAST (v2.7.1)^129^ was used to search for L5A5 repeats elsewhere in the HLE assembly.

To localize the L5A5 repeat on HLE metaphase chromosome spreads, a 33 bp oligo was designed specific to the L5A5 repeat. The oligo was diluted to a concentration of 100 μM, then a 3’ end labeling reaction was set up according to the following: 4 uL 5x TdT reaction buffer, 4 uL 25 mM CoCl_2_, 1 uL 1 mM dig-dUTPs, 1 uL TdT transferase enzyme, 6 uL ddATPs, and 3 uL water (20 uL total reaction). The reaction was incubated at 37°C for 30 minutes, then the reaction was stopped using 2 uL 0.2 mM EDTA. DNA FISH was performed on HLE metaphase chromosome spreads as published previously^10^. Briefly, prior to slide preparation for hybridization, slides were dehydrated in 100% EtOH, then air dried. Slides were treated with 200 uL 0.1mg/mL RNase A in 2X SSC and incubated at 37°C for 15 minutes in a humid chamber with a parafilm coverslip, then rinsed 4 times with 2X SSC pH 7.0 for 2 minutes each. Following, slides were treated with HCl (49.2 mL water + 0.8 mL 12N HCl) for 10 minutes, then rinsed in 2X SSC pH 7.0. Lastly, slides were dehydrated in a 70%, 90%, and 100% EtOH row for 2 minutes each before air drying. To denature the slides, they were treated with 70% formamide in 2X SSC at 72°C for 2 minutes, then transferred to a −20°C 70%, 90%, and 100% EtOH row for 2 minutes each before air drying. Meanwhile, 5 uL of the L5A5 oligo probe was precipitated with 1 uL of 10 ug/uL salmon sperm carrier DNA and 2.5X volume of 100% cold EtOH at −80°C for 40 minutes. After precipitation, the probe was centrifuged at 13,000 rpm for 25 minutes. The supernatant was removed, and the probe was resuspended in 12 uL Hybrisol VII (MP Biomedicals). The probe was rehydrated for 1 hour, then denatured at 80°C for 5 minutes. The probe was applied to the slide, then incubated overnight with a sealed coverslip at 37°C in a humid chamber. The following day, the coverslip was removed, and slides were washed 1 time for 2 minutes at 72°C in 0.4X SSC/0.3% NP-40, then 1 time for 1 minute at room temperature in 2XSSC/0.1% NP-40. Subsequently, the slides were rinsed on 0.2% Tween 20/4XSSC, then blocked for 30 minutes at 37°C in 0.2% Tween 20/4XSSC/5% BSA. Slides were rinsed in 0.2% Tween 20/4XSSC, then an anti-Dig fluorophore was applied at a 1:400 dilution at 37°C for 30 minutes. Slides were rinsed three times in 0.2% Tween 20/4XSSC for 5 minutes each at 45°C, rinsed in H_2_O, then serially dehydrated in a 70%, 90%, and 100% EtOH row for 2 minutes each. After air drying, a counterstain of DAPI (diluted 1:5 in Vectashield) was applied to the slides and covered with a coverglass. Slides were imaged on an Olympus AX70 microscope using CytoVision software (Leica Biosystems Richmond, Inc.).

Chromosome painting was carried out on HLE metaphase chromosome spreads using Aquarius Whole Chromosome Painting probes (Cytocell Ltd) for human chromosome 20. Slides and probes were co-denatured at 72°C for 5 minutes on a Hybaid in-situ block, then probes were hybridized to slides treated above in a humid chamber overnight at 37°C. Post hybridization, slides were washed in 2XSSC at room temperature to remove the coverslip, then washed in 0.4XSSC at 60°C for 2 minutes and 2XSSC/0.5% Tween 20 for 1 minute at room temperature. Slides were then rinsed in distilled water, dehydrated in a 70%, 90%, and 100% EtOH row, then counterstained as above with a 1:5 dilution of DAPI in Vectashield (Vector Laboratories, Inc.). As above, images were captured using an Olympus AX70 microscope and CytoVision software (Leica Biosystems Richmond, Inc.).

### Copy number estimation of L5A5 and evaluation of L5A5 repeats across primates

To estimate the copy number of L5A5 repeats in the HLE genome, a *k*-mer based approach was taken as developed for the rDNA copy number estimation in T2T-CHM13 genome assembly^11^. First, 21-mers in the HLE PCR-Free Illumina reads were counted using Meryl (v1.3)^18^, then 21-mer multiplicity (or number of times the 21-mer was found in the read set) was plotted against the corresponding counts of 21-mers found at that multiplicity. The average 21-mer multiplicity was determined to be 46, corresponding to an average sequencing depth of 46X. Next, the coverage of 21-mers in the Illumina PCR-Free read set was determined across the L5A5 consensus sequence using Meryl v(1.3)^18^. The corresponding counts were divided by 23X, the estimated read coverage per haplotype, to determine the diploid copy number of L5A5 repeats. The median was determined to be ∼1,154 total copies in the genome, or ∼577 copies per haplotype.

In order to assess the presence of L5A5 repeats across other primates, BLASTN (v2.7.1)^129^ was used to search for L5A5 composite repeats across 13 additional primates. The start and stop locations of identified repeats were curated using the Geneious software (build 2021-07-19 12:20)^128^. The sequences were aligned to the HLE consensus using a Geneious alignment with default high sensitivity settings.

### L5A5 copy number validation with PCR

To validate the L5A5 repeat array identified in HLE and not in other species, primers were designed to target a 774 bp internal portion of L5A5 repeats (expected to amplify in both HLE and NLE) and a 774 bp portion of the junction between two L5A5 repeats (expected to amplify in only HLE). A PCR reaction was set up according to the following: 1 uL 10 uM F primer, 1 uL 10 uM R primer, 1 uL Taq polymerase, 2.5 uL 2.5 mM dNTPs, 5 uL 10X PCR buffer, 120 ng of HLE or NLE DNA, and H_2_O to 50 uL. Reactions were cycled in a thermal cycler according to the following: 94°C for 3 minutes, then 94°C for 30 seconds, 58°C for 30 seconds, 72°C for 1 minute (repeated for 35 cycles), then 72°C for 5 minutes. Amplification was visualized on a 1% agarose/1X TBE gel with EtBr, run at 90V for 50 minutes. The gel was imaged on a BIO RAD GelDoc™ EZ Imager using the Image Lab (v6.0.1) software.

### Evaluation of breakpoints in the HLE assembly

Breakpoints corresponding to T2T-CHM13, NLE, HMO, and SSY were identified in the HLE assembly by performing pairwise alignments between the HLE assembly and each other genome using LASTZ (v1.04.15)^101^. Filtering and chaining was performed with UCSC GenomeBrowser Kent Tools (v369)^102^. A custom python script, axtToSyn^130^, was used to detect synteny blocks and breakpoints using a minimum alignment length of 1,000 bp and a minimum alignment score of 100,000. Identified breakpoints, reported by axtToSyn as the 1,000 bp flanking the left and right of breakpoints, were collapsed into one set of coordinates. Breakpoints and syntenic blocks were plotted using RIdeogram^131^. Identified breakpoints were intersected with RepeatMasker annotations using Bedtools (v2.29.0) intersect and summarized using the RepeatMasker utility script buildSummary.pl^109^.

To analyze methylation of repeats at breakpoints, intersected repeats were filtered for one of: L1Hylob, L1Ms, L1P (LINEs), AluJ, AluS, AluY (SINEs), or LAVAs. Repeats found within each breakpoint were concatenated together, then Bedtools (v2.29.0) subtract^115^ was used to remove TEs found within breakpoints from genome wide TEs, to create five categories: breakpoints to T2T-CHM13, breakpoints to HMO, breakpoints to NLE, breakpoints to SSY, and TEs found outside of breakpoints. Methylartist (v1.2.7) segmeth^56^ was used to output aggregated CpG methylation calls over intervals for each repeat. Following, Methylartist (v1.2.7)^56^ segplot was used to create ridge plots for each repeat type. Aggregated methylation calls over intervals as called by Methylartist (v1.2.7) segmeth^56^ for each repeat type were averaged to get average methylation per repeat type per breakpoint category. Descriptive statistics were calculated using GraphPad Prism (v9).

To analyze segmental duplication content of breakpoint regions, SD calls from *BISER* (v1.4)^40^ were intersected with breakpoints using Bedtools (v2.29.0) intersect^115^. SD intersect statistics were calculated using GraphPad Prism (v9). To identify older (HyA) and young (HLE) breakpoints, HLE-CHM13 breakpoints corresponding to those between syntenic blocks identified in ^2^ were subsetted; other breakpoints were excluded from the analysis due to lack of age evidence. Subsetted breakpoints were aged by identifying those present in the gibbon ancestor (HyA) or not found in the gibbon ancestor (HLE). Comparisons between aged breakpoints were made by intersecting varying data types with breakpoints using Bedtools (v2.29.0) intersect^115^ and statistical comparisons were performed using the Mann-Whitney U test.

### Primary processing of Omni-C data

Raw Omni-C data was aligned to the reference HLE genome using bwa (v7.0.17)^90^ with the following settings (mem -5SP -T0 -t16). Following alignment, the parse module of pairtools (v1.0.2)^96^ was utilized (*--min-mapq 40 --walks-policy 5unique --max-inter-align-gap 30*) to identify ligation junctions and outputs were sorted using the pairtools sort module with default settings. Next, the dedup module was used to detect and remove PCR duplicates. To generate bam files for downstream analysis, we used pairtools to split, sort and index the .pairsam file with default settings. Using HiCRes^132^ we estimated resolution of the Omni-C dataset and used the cooler tool v0.9.3^133^ to convert the indexed output file into a single resolution cool matrix (10kb bin size) and the multi-resolution mcool matrix (resolutions ranging 10kb-10.24Mb). Output files were compressed using samtools bgzipm^134^ . TAD boundary annotations and genome-wide insulation scores were generated by HiCExplorer (v3.7.2) ^135^ with a 10Kb resolution (--minDepth 100000 -- maxDepth 600000).

### Genome conformation and CTCF binding motif annotation at evolutionary breakpoints

We used previously published^52^ custom scripts (https://github.com/carbonelab/hicpileup) to visualize median insulation score in 600 Mb windows centered at the synteny breakpoints identified against CHM13, NLE, HMO and SSY. The same scripts were used to visualize aggregate Hi-C contact frequencies in 2Mb windows centered at synteny breakpoints. To further explore the relationship between chromatin interaction and age of breakpoints, for each breakpoint we determined minimum and median insulation score as well as distance from the closest TAD boundary, using bedtools and custom shell and R scripts. Bedtools shuffle (v2.31.1) (-chrom - noOverlapping) was used to generate a shuffled version of each set of breakpoints followed by two-tailed wilcoxon rank-sum tests to compare distance to nearest boundary, and minimum/median insulation score between observed and shuffled breakpoints. We used the findMotifsGenome.pl (size -given) function from Homer (v4.11)^136^ to predict all CTCF binding motifs within sequences corresponding to all BOS sites.

### Associations between genomic features and nucleotide diversity

All paired-end illumina data were retrieved from the NCBI database for *Hoolock leuconedys* using fastq-dump from the sratoolkit (v.3.0.0)^137^ (accessions: SRR10075429, SRR10075430, SRR10075432, SRR10075433), and adapters were then trimmed from paired-ends using fastp v.0.23.4^138^.

The genome assembly fasta was indexed using bwa index^139^, followed by alignment and sorting each sample using bwa mem with the Sentieon workflow under default parameters. We then applied the LocusCollector algorithm from Sentieon to the bam alignments to collect read information and applied Sentieon’s Dedup algorithm to mark and remove PCR duplicates. Next, we used Sentieon’s Haplotyper algorithm, using emit_mode gvcf and default filters, to produce a gvcf file for each sample. This was followed by joint genotyping using the GVCFTyper algorithm, under default filtering settings.

We then generated a mappability filter using GenMap v.1.3.0 and retained only sites that map to the genome uniquely with a kmer size of 150^140^. In addition to this filter, we removed sites with map quality < 40 or quality scores < 20 using bcftools v.1.20^134^. We then assessed sample depth, missingness, and heterozygosity using vcftools v.0.1.16, and output the average nucleotide diversity (π) with window sizes of 100 kb^141^. Using GFFUtils v.0.13^142^, we converted the GTF annotation file into bed format, and used a series of custom scripts to calculate the distance to the nearest breakpoint and the centromere from the midpoint of each window. We also calculate the percentage of each window covered by an annotated gene,segmental duplications, repeat content, as well as SST1, LAVA, L1P, L1M, L1Hb, *Alu*Y, *Alu*S, *Alu*J, and *Alu*Jb elements specifically.

In R^143^ we performed a linear model on the above described data as predictors, for the following interaction effects: distance to the nearest breakpoint*breakpoint age, minimum insulation score of the nearest TAD boundary* the closest TAD boundary distance* the distance to the nearest breakpoint, with nucleotide diversity as the response variable. Due to lack of normality in the residuals, we performed an ordered normalization transformation on the nucleotide diversity values with the bestNormalize package^144^. We then use the backwards step function from the car package^145^ to determine the model with the best fit and evaluate interaction effects using the plot_model function, with type = “eff”, from the sJplot package^146^.

## Supplementary Information

### Supplemental Tables

**Table S1: Repeats across the HLE assembly.** Repeat counts, basepairs, and corresponding basepair percentage of the HLE assembly are listed for major repeat classes as defined by RepeatMasker.

**Table S2. Span of CENP-A enrichment across assembled HLE chromosomes.** The start, end, and total length in basepairs of CENP-A CUT&RUN mapping enrichment is denoted for each chromosome. Evidence of collapses (YES or NO) for each centromere as defined by spikes in read coverage is included.

**Table S3: Repeat content of each assembled HLE centromere.** The percentage of CENP-A enrichment spans that correspond to annotations for each of the major repeat classes by basepair defined by RepeatMasker. The total repeat content of the CENP-A enrichment spans are aggregated below.

**Table S4: Genes present within 200kb of assembled HLE centromeres.** The chromosome, start, stop, gene orientation, FLAG gene ID, Eggnog ortholog, gene name, Eggnog taxscope, and gene drescription as reported by FLAG are reported for genes within 200kb of CENP-A enrichment.

**Table S5: Top ten GO biological processes enriched among genes within 200kb of assembled HLE centromeres.** The p-value, odds ratio, and combined score are reported.

**Table S6: Consensus sequence of L5A5.**

**Table S7: Oligo sequence for probe targeting the L5A5 repeat.**

**Table S8: Species searched for the L5A5 repeat.** The representative phylogenetic grouping, scientific name, common name, and accession numbers are shown for primates included in Figure 4E.

**Table S9: Primer information for primer sets targeting the L5A5 repeat.** Primers target both internally (within a single L5A5 repeat) and at the junction (between two L5A5 repeats).

**Table S10: Breakpoint locations across the HLE genome.** The table reflects a bed file of all breakpoint locations, with a fourth column representing which genome the breakpoint is respective to.

**Table S11: Breakpoint locations across the HLE genome.** The table lists the chromosome, start, and stop location of all breakpoints, collapsed to show those shared respective to multiple genomes. Columns indicate whether a breakpoint is found respective to another genome (CHM13, HMO, NLE, or SSY) or at a centromere (HLE_Chr_# indicating which centromere). If not shared with a particular genome or centromere, the breakpoint is indicated by a . in that column.

**Table S12: Percentage of repeats overlapping with HLE breakpoints respective to T2T-CHM13, NLE, HMO, and SSY.** Repeat percentages are listed as a percentage of all annotated repeats in each corresponding category. Overall repeat percentages are included for comparison.

**Table S13: Aggregated CpG methylation statistics for SINEs (AluY, AluS, AluJ), LINEs (L1Hylob, L1M, L1P), and LAVAs.** The number of each corresponding repeat type containing methylation calls overlapping with breakpoints and the minimum, median, mean, maximum, and coefficient of variation of CpG methylation across repeats is listed for breakpoints respective to T2T-CHM13, HMO, NLE, and SSY. For comparison, statistics are included for repeats in each category not found in any breakpoint regions. The difference in medians is included between each breakpoint category compared to repeats found in non-breaks, and cells are color coded from green (repeats are more methylated at breaks) to red (repeats are less methylated at breaks). Mann-Whitney p values comparing the two groups are included in the final column.

**Table S14: Number of breakpoints associated with segmental duplications.** The number of HLE breakpoints respective to T2T-CHM13, HMO, NLE, and SSY are listed, with corresponding counts of breakpoints overlapping with segmental duplications and their associated average basepair coverage.

**Table S15: Chromatin conformation statistics at HLE breakpoints (BOS) respective to T2T-CHM13, HMO, SSY and NLE.** In each species we find BOS to be located significantly closer to TAD boundaries and display smaller minimum/median insulation scores, compared to random expectation (i.e., shuffled BOS).

**Table S16: Statistics for a linear model of interaction effects are shown for various coefficients with nucleotide diversity as a response variable are shown.** The table includes standard error, the t value, p value, and significance. A negative estimate indicates decreased diversity, and vice versa. A key for easier interpretation is included. See Figure S14 and S15.

### Supplemental Figures

**Figure S1: RNA polymerase occupancy derived from PRO-seq alignments conforms with expectations at both gene promoters and bodies across the genome.**

**(A)** A principal component analysis (PCA) plot shows a genome-wide view of PRO-seq alignment replicates. PRO-seq replicates (mapped with Bowtie2 (default, “best match”))^123^ are highly similar based on a Spearman Correlation with a coefficient of 0.91. Scatterplot shown as the natural log after adding 1 with outliers removed. Below, heatmaps display RNA polymerase position per gene (horizontal lines) for antisense (blue) and sense (red) strands based on Bowtie2 mapping (default, “best match”)^123^, while composite profiles (along the top) display a summary of all genes collectively. **(B)** The heatmap highlights the expected RNA polymerase accumulation at gene promoters. Each gene is anchored at its 5’ end (transcription start site; TSS) with distances indicated in the bottom right (5.0kb into the gene body) and bottom left (1.0kb away from the gene body). The dotted line marks the fixed end of each gene, with genes ordered from longest to shortest, top to bottom. **(C)** The heatmap highlights the expected RNA polymerase presence across the entire gene body: enriched at the promoter and lessened across the body, with a modest increase at the 3’ end (demonstrating a slowdown of polymerase activity as termination progresses). Each gene is scaled to the same length with distances on either side of the gene body indicated in the bottom corners (0.1kb). Genes are ordered based on transcriptional signal intensity from strongest to weakest, top to bottom.

**Figure S2: Transcriptional profile of repeats based on PRO-seq.**

Distribution of the 16 repeat classes (excluding rRNA), quantified by either Bowtie or Bowtie2^123^ using three different settings for handling multi-mappers: (shown from left to right) Bowtie2 default (single alignment per read, “best match”), Bowtie k-100 (up to 100 alignments allowed per read), and Bowtie k-100 21-mer (k-100 option subsequently filtered through genome-wide single-copy 21-mers).

**Figure S3: NucFreq**^50^ **long-read read coverage analysis of assembled HLE centromeres confirms assembly of six HLE centromeres.**

Read coverage analysis of six assembled HLE centromeres (Cen1, Cen3, Cen8, Cen9, Cen11, and CenX) shows generally even coverage across the span of CENP-A and the surrounding 100 kilobases, suggesting successful assembly of the centromeres, with fluctuations in read coverage (black and red data points) suggesting only minor nucleotide errors. Below read coverage maps, CENP-A enrichment is depicted as CENP-A CUT&RUN reads mapping that overlap with unique 21-mers to provide a conservative estimate of CENP-A localization (blue). Repeat annotations are depicted along the region, with each major class of repeats represented by a different color according to the key below. Average CpG methylation is represented by a line plot.

**Figure S4: NucFreq**^50^ **long-read read coverage analysis of assembled HLE centromeres confirms assembly collapse of 13 HLE centromeres.**

Read coverage analysis of 13 assembled HLE centromeres reveals large spikes in mapped read coverage (black and red data points), suggesting the assembly is collapsed in those regions.

Centromeres are shown as the estimated span of CENP-A enrichment plus 100 kilobases on either side. As described in Figure S3, additional tracks show CENP-A enrichment, repeats, and CpG methylation across the loci.

**Figure S5: Centromere assemblies of Chr1, Chr3, and Chr8.**

The centromeric region of HLE chromosome 1 **(A),** HLE chromosome 3 **(B)**, and HLE chromosome 8 **(C)** are shown. Per Figure 2, from top to bottom, genome tracks denote CENP-A CUT&RUN enrichment (blue), repeat annotations colored according to the key, synteny to T2T-CHM13, and predicted HLE-CHM13 synteny breakpoints. Fiber-seq inferred regulatory element (FIRE) tracks show FIRE density binned per 1kb on a heatmap scale from white to black (i.e. low to high density), showing increased density of FIREs correlating with CENP-A enrichment corresponding to dichromatin organization. Gene tracks show gene predictions from FLAG. Replication timing from E/L Repli-seq is shown as black points indicating the log ratio of early-to-late coverage over 5kb windows from 4 (early replication) to −4 (late replication), with a red line indicating the 10 point moving average. CpG methylation is shown via line plot (black line) and on a heatmap scale from low CpG methylation (black) to high CpG methylation (red), and CG density is shown (purple).

Finally, sequence identity plots are shown for each assembled centromere, with a scale from blue (low identity) to red (high identity). **(D)** The panel shows a dot-plot (Gepard^82^, word length 50) comparing HLE Cen3 (HLE_Chr_3:74500882-74510287) to the SSY Cen1 (mSymSyn1_v2.0 chr1_hap1:81931844-82305723) recently released^127^. Corresponding alpha satellite annotation tracks for both centromeres show alpha satellite super families (SFs) and the strand orientation according to the key. The SSY Cen1 is much larger (∼370 kb), formed entirely by SF4 (yellow). A smaller (∼9 kb) alpha satellite of the HLE centromere corresponds to the leftmost part of the siamang array as indicated by the dot-plot diagonal (identity ∼98%). Within the SF4 (yellow) arrays, light blue stripes indicate occasional presence of R2 (SF5) monomers in the arrays mostly formed by Ga (SF4) monomers. These 2 monomeric types are very similar in sequence, with only 4 differences between the consensus sequences, and can easily be misclassed into each other in divergent monomeric arrays. Such contamination is common and does not change the classification of the arrays as SF4^8^.

**Figure S6: Unlike highly homogenized human centromere cores, the CENP-A binding domain is not a transcriptional dead zone.**

Plots for Cen 1 **(A)**, Cen3 **(B)**, Cen8 **(C)**, Cen9 **(D)**, Cen11 **(E)**, and CenX **(F)** show CENP-A enrichment (blue), repeat annotations represented by one color per repeat category, and HLE RNA-seq mapped across the region. Above and below the repeat track, PRO-seq tracks show RNA polymerase activity along the positive and negative strands at three-levels of stringency (from least to most): Bowtie K100 (grey), Bowtie K100 21-mer filtered (red), and Bowtie2 default (black). Below, CpG methylation (black line) shown additionally as profiles generated by Methylartist^56^. Each methylation profile panel includes methylated CpGs denoted as teal and unmethylated CpGs denoted as black. Below, m6A methylation from Fiber-seq is shown (blue).

**Figure S7: HLE Cen8 CENP-A enrichment overlaps with clustered mitochondria homolog pseudogenes (*CLUHP)*.**

Overall sequence similarity between HLE *CLUHP* (5’) in the middle of HLE Cen8 and HSA *CLUHP* and HLE *CLUHP* (3’) genes are shown. If more than one *CLUHP* pseudogene was found on the same chromosome, the genes are denoted by their position relative to one another, 5’ or 3’. Apart from the 3’ HLE ortholog sharing 91% sequence similarity, HLE *CLUHP* 5’ shares 80-90% sequence identity with all identified HSA pseudogenes. **(B)** Two full length copies of *CLUHP* were identified in the Cen8 region **(Figure S5C)**, each containing eight exons. Sequence alignment between each exons is shown, with each nucleotide highlighted green if at least 70% of the aligned sequences have the same DNA base as the reference sequence, HLE *CLUHP* 5’.

**Figure S8: The *PLSCR* gene family is disrupted by evolutionary breakpoints in HLE, yet *PLSCR1* maintains expression.**

**(A)** From top to bottom, genome tracks denote CENP-A CUT&RUN enrichment (blue heat map), repeat annotations colored according to the key, CpG methylation, synteny to T2T-CHM13, and gene annotations for HLE Cen9 (top left panel) and HLE Chr 6 (top right panel), showing the breakpoint regions involved in the *PLSCR* gene family rearrangement separating *PLSCR4* to HLE Chr 9 and *PLSCR5* and *PLSCR1* on HLE Chr 6. The full length *PLSCR2* was not detected; however, a duplicated portion of the gene is present. Below gene annotations, tracks show RNA-seq mapping for HLE, NLE (Vok), and PRO-seq Bowtie2 default mapping. Below HLE panels, the structure of the *PLSCR* gene family in humans is shown, with repeat and gene tracks showing the homologous regions of T2T-CHM13 on HSA Chr 3 and HSA Chr 12. Using T2T-CHM13 RNA-seq and PRO-seq^21^, the transcriptional status is classified below gene names by evidence of transcription in T2T-CHM13 (green circles), some evidence of transcription in T2T-CHM13 (ie., 1-2 exons show RNA/PRO-seq signal, orange circles), or no evidence of transcription (red circles). Unlike T2T-CHM13, most genes in the rearrangement regions appear transcriptionally inactive in HLE; however, this difference may be cell type dependent. However, *PLSCR1* still shows high levels of transcription in HLE. **(B)** A detailed view of the *PLSCR* breakpoint organization is shown in humans (T2T-CHM13, Chr 3), rhesus macaque (Mmul_10/rheMac10, Chr 2), and common marmoset (Callithrix_jacchus_cj1700_1.1/calJac4, Chr 17), not to scale. In all three lineages, *PLSCR1* (nine exons), *PLSCR2* (eight exons), and *PLSCR2-*like (six exons) are present. Exons are indicated by dark blue, light blue, dark purple, and light purple, with duplicated exons shared by *PLSCR1* and *PLSCR2* indicated by light blue. Predicted exons that may be misannotations (ie., TEs) are indicated by red highlighted boxes. Teal boxes indicate sequences in between genes or repeat-rich regions, and breakpoints are indicated by red stars. In HLE, this organization was disrupted by a chromosome rearrangement, with *PLSCR1* and *PLSCR2* (exon one only) found on HLE Chr 6, and *PLSCR2*-like found on HLE Chr 9. The full length *PLSCR2* was not detected in HLE.

**Figure S9: Segmental duplications of SST1 and LAVA elements are present in pericentromeres.**

**(A)** A circos plot representing the 19 assembled HLE chromosomes is depicted. As in Figure 1C, each chromosome is marked in a different color, and is represented by the outer track with ticks along each chromosome representing 10 megabases. Segmental duplications filtered on length (>1 kb) and identity (>90%) are shown as linkages between chromosomes. **(B)** As in **(A)**, the 19 assembled HLE chromosomes are depicted. Predicted centromere regions are marked in track 2 by black lines. Filtered segmental duplications overlapping LAVA annotations are denoted by red linkages, and filtered segmental duplications overlapping SST1 annotations are denoted by purple linkages. We speculate that these large inter-chromosomal segmental duplications, often impacting the centromere or surrounding pericentromere, may help facilitate the rapid karyotype evolution of gibbons.

**Figure S10: The HLE latent Cen17 shares similarity to the flanks of SSY Cen24.**

The panel shows a dot-plot (Gepard^82^, word length 50) comparing the inactive HLE Cen17 alpha satellite array (HLE_Chr_17:38245650-38275051) to the two flanks of the recently released SSY Cen24 (mSymSyn1_v2.0 chr24_hap1:46819441-46910555, chr24_hap1:50119536-50210782)^127^. Corresponding alpha satellite annotation tracks for both centromeres show alpha satellite super families (SFs) and the strand orientation (red and blue). The dot-plots suggest that the most of the latent HLE Cen 17 corresponds to the rightmost part of the SSY Cen24 array with some deletions in SSY indicated by breaks in the diagonal (lower close-up, identity ∼98%). A small portion of alpha satellite at the left tip of the latent HLE Cen17 array corresponds to a portion of the SSY gibbon Cen24 array on the left flank (upper close-up, green frame, identity ∼97%). Thus, pieces of both flanks of an HLE-SSY ancestral centromere, which likely looked similar to the current SSY Cen24 array, are joined in the HLE array. This site marks the old centromere location, while the current HLE centromere has shifted to another location. Additionally, the yellow (SF4) arrays have intermittent light blue stripes indicating occasional presence of R2 (SF5) monomers in the arrays, mostly formed by Ga (SF4) monomers. These two monomeric types are very close to each other, with only 4 differences between the consensus sequences, and can easily be misclassed into each other in divergent monomeric arrays. Such contamination is common and does not change the classification of the arrays as SF4^8^.

**Figure S11: NucFreq**^50^ **analysis of a latent centromere on Chr17 confirms successful assembly free from collapses.**

On top, a panel shows NucFreq long read coverage analysis of the latent HLE Cen 17 region, confirming that there are no assembly collapses in the region despite the small array size. Below, a CpG methylation track shows the lack of a CDR, repeat annotations, and CENP-A coverage indicating the region no longer binds CENP-A.

**Figure S12: PCR of L5A5 confirms the repeat is arrayed in HLE, but not in other gibbons (NLE).**

To confirm the computational observation that L5A5 is arrayed in HLE and not in other gibbons, two PCR primer sets were designed targeting L5A5: first, a set was designed to target the junction of two L5A5 composite monomers in an array, expected to amplify a 774 bp product only in HLE; and second, a set was designed to target the L5A5 monomer internally, expected to amplify a 774 bp product in HLE *and* other species, with a higher degree of amplification in HLE given the repeated structure (Table S9). When tested on closely related gibbon species *Nomascus leucogenys* (NLE), the primer sets worked as predicted, amplifying a product only in HLE when targeting the junction between two monomers and amplifying a product both in HLE and NLE when targeting the internal structure, albeit at a lower concentration in NLE. These results indicate that the L5A5 repeat is arrayed in HLE, but not in NLE.

**Figure S13: CpG methylation status of *Alu*Y, *Alu*J, *Alu*S, L1M, and L1P repeats within or outside of breakpoints.**

As in Figure 5D, aggregated CpG methylation across *Alu*Y, *Alu*S, *Alu*J, L1P, and L1M repeats are depicted as ridgeplots, showing repeats annotated within BOS respective to CHM13, HMO, NLE, and SSY, as well as repeats outside BOS. Significance for comparisons between methylation at breakpoints and outside of breakpoints can be found in Table S13.

**Figure S14. Predicted values of nucleotide diversity as a function of the interaction between distance to the nearest chromosomal breakpoint and the closest TAD boundary.**

The X-axis represents the distance to the nearest chromosomal breakpoint in base pairs (bp), and the Y-axis represents nucleotide diversity, with the data stratified by the distance to the closest TAD boundary. Nucleotide diversity generally increases with greater distance from the nearest chromosomal breakpoint, with the effect varying depending on the closest TAD boundary.

**Figure S15. Predicted values of nucleotide diversity as a function of the interaction between breakpoint age and distance to the nearest breakpoint.**

The Y-axis represents nucleotide diversity, and the colors represent different distances to the nearest chromosomal breakpoint. Nucleotide diversity shows variation with breakpoint age, and this relationship is influenced by the distance to the nearest chromosomal breakpoint, with greater distances generally associated with higher diversity values, but to a higher degree in association with older breakpoints.

**Figure S16. With the exception of insulation scores and presence of LAVA, no other genomic or epigenetic feature distinguishes HyA versus HLE breakpoints.**

As in Figure 5G, dot plots show differences, or lack thereof, between older BOS (HyA) and younger (HLE) BOS. Statistical comparisons were performed using the Mann-Whitney test.

