## Supplementary Figures for "Centromeric transposable elements and epigenetic status drive karyotypic variation in the eastern hoolock gibbon"

**Hartley, et al _ Figure S1**

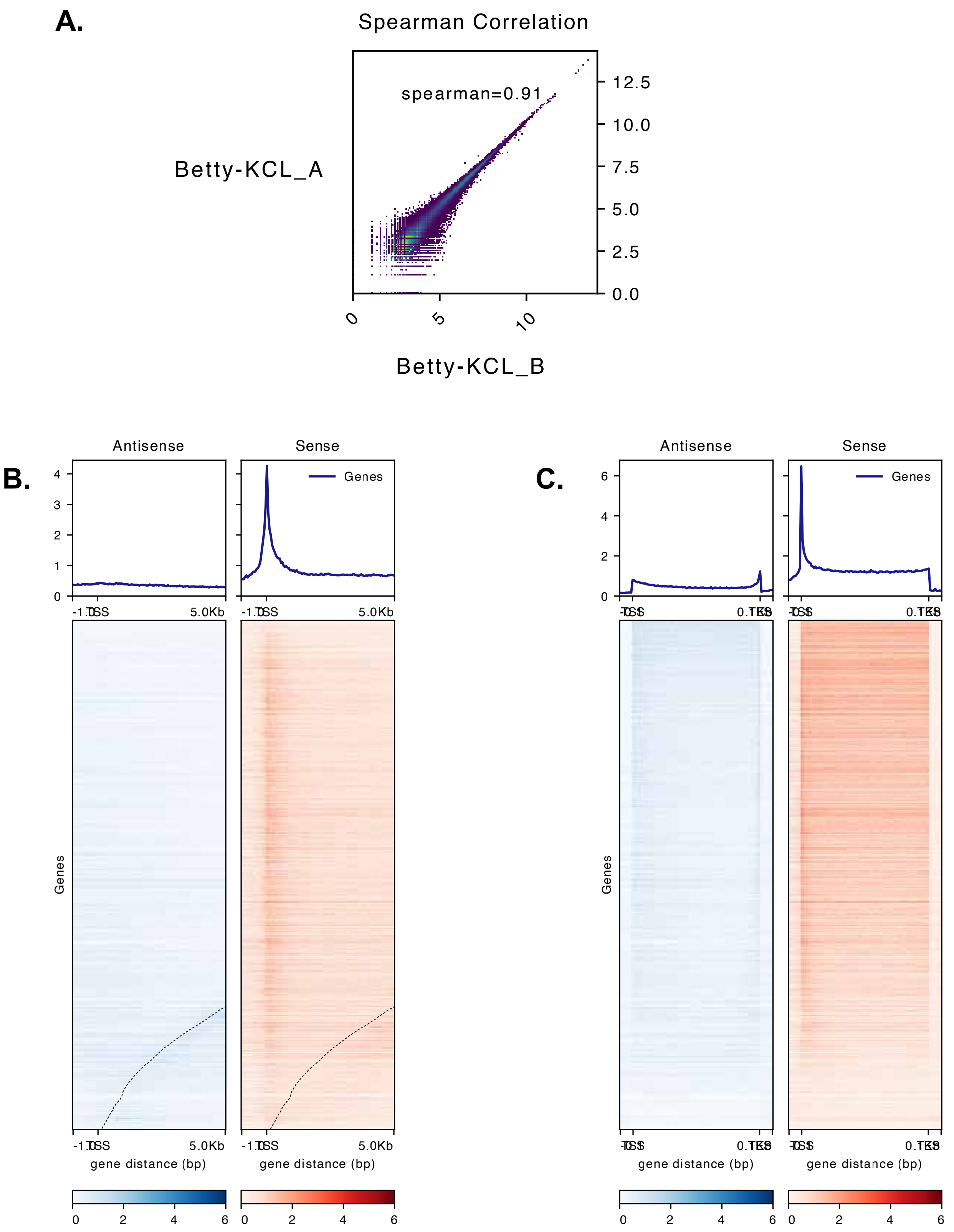

Hartley, et al _ Figure S2

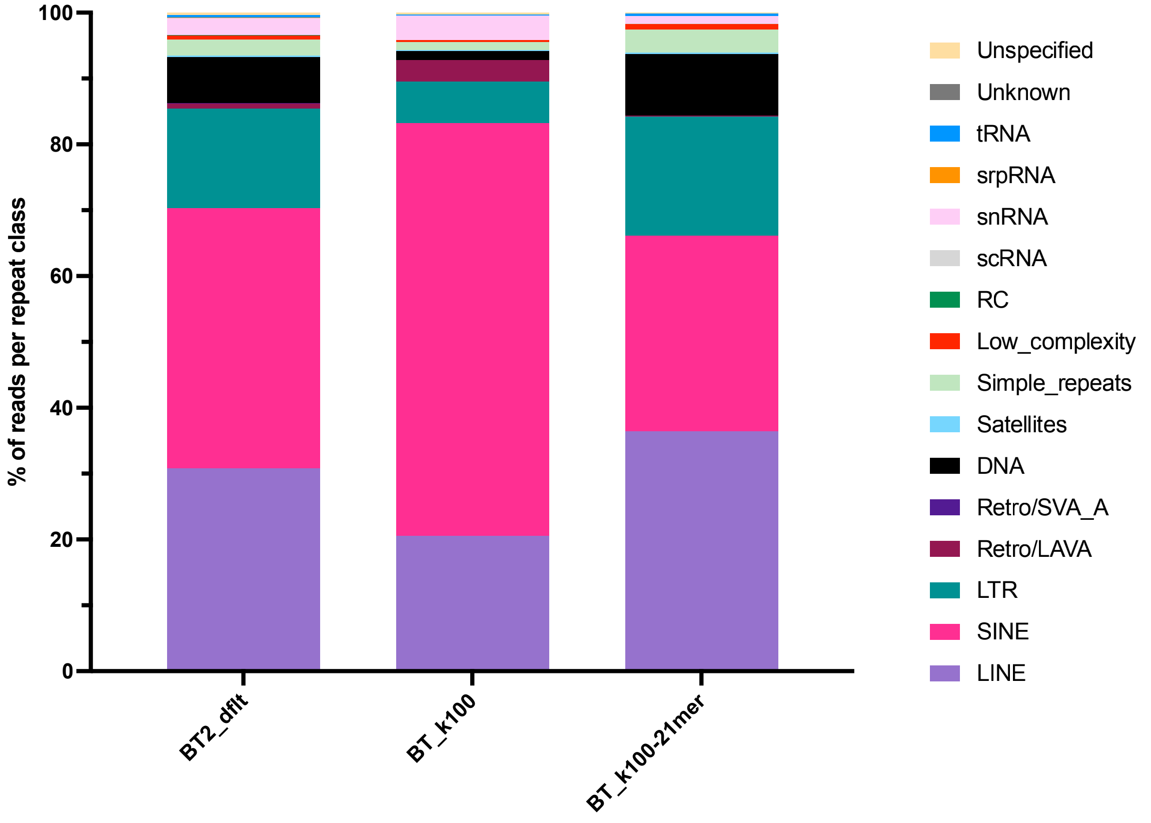

### Hartley, et al _ Figure S3

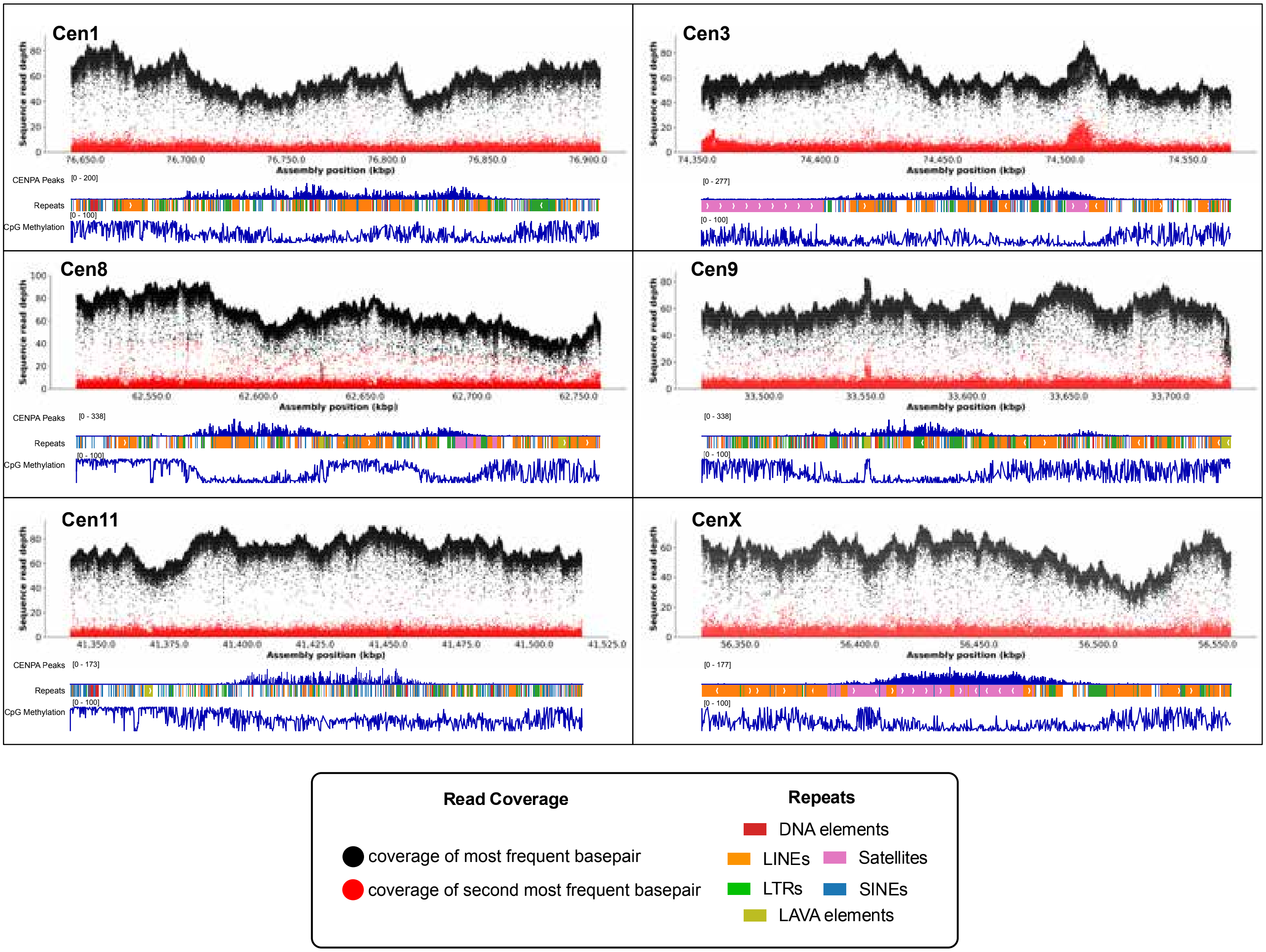

### Hartley, et al _ Figure S4

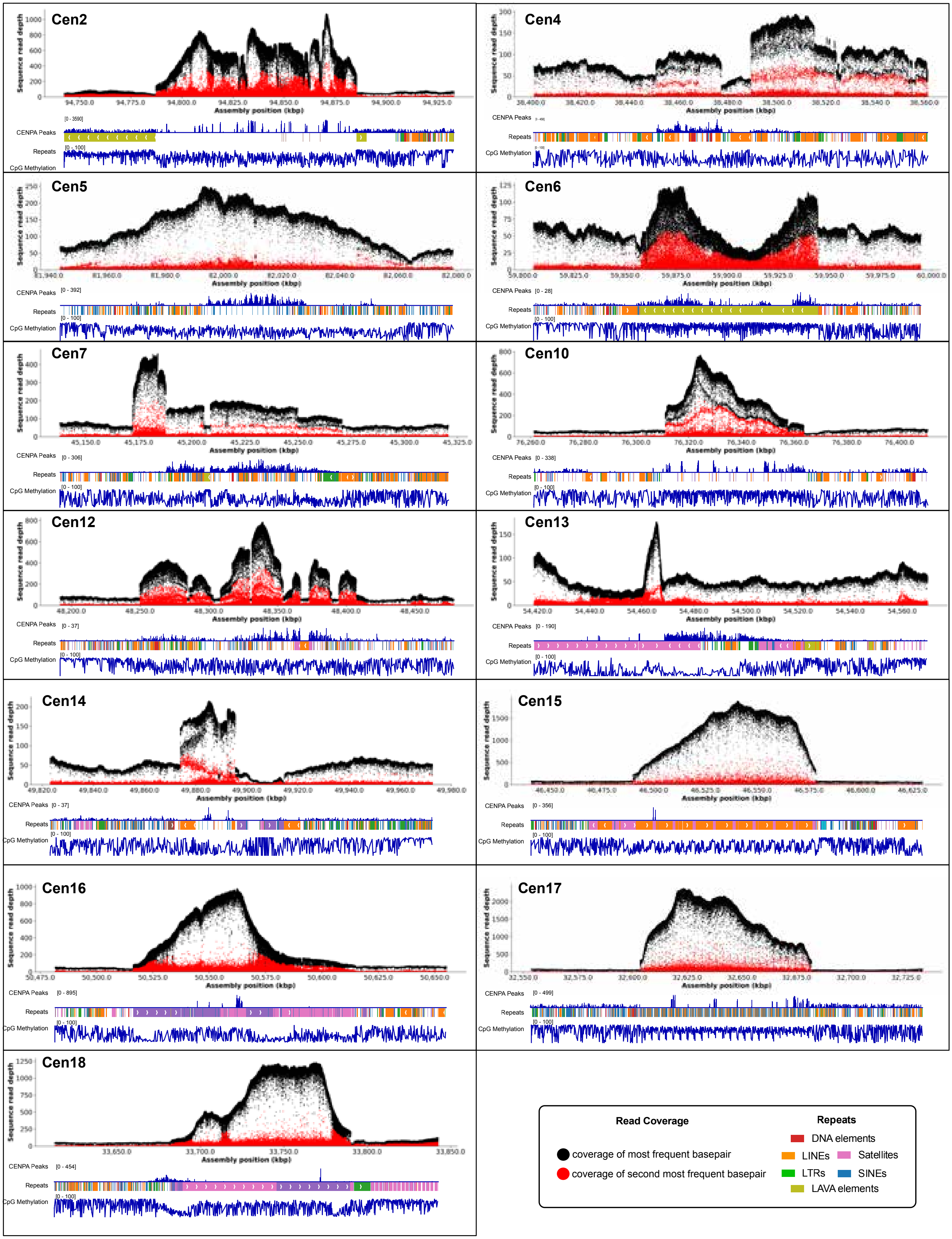

Hartley, et al _ Figure S5

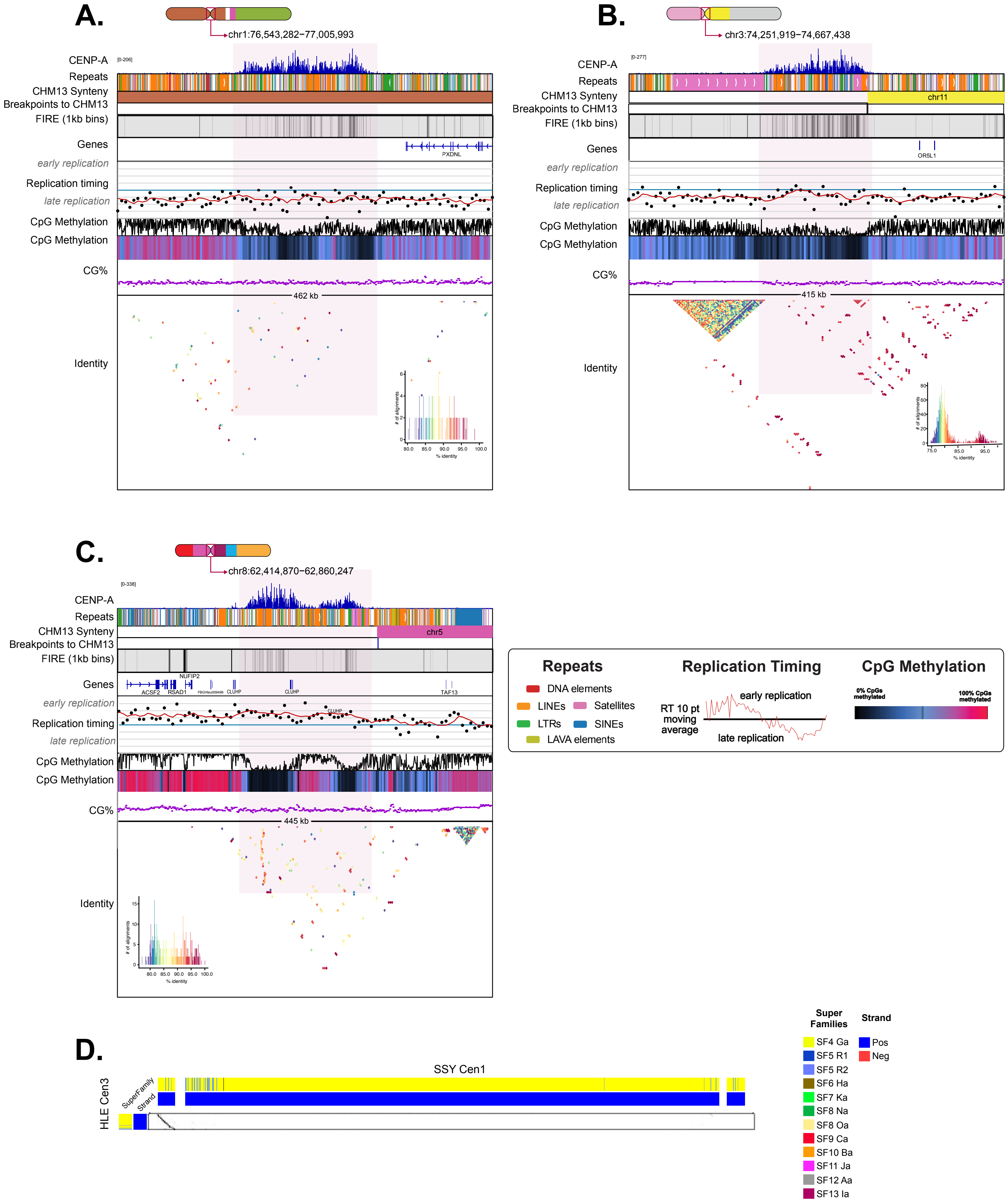

### Hartley et al, Figure S6

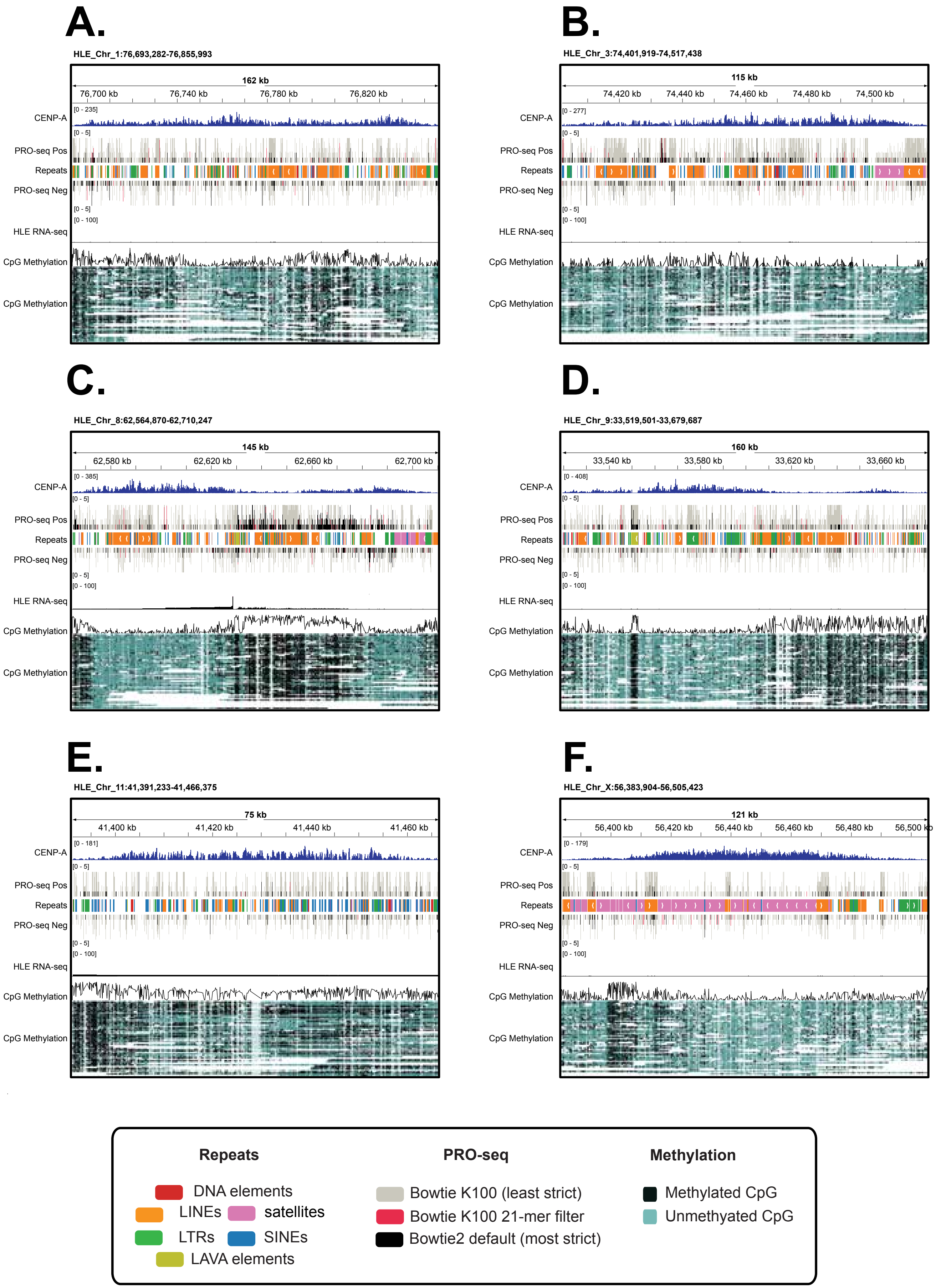

### Hartley et al, Figure S7

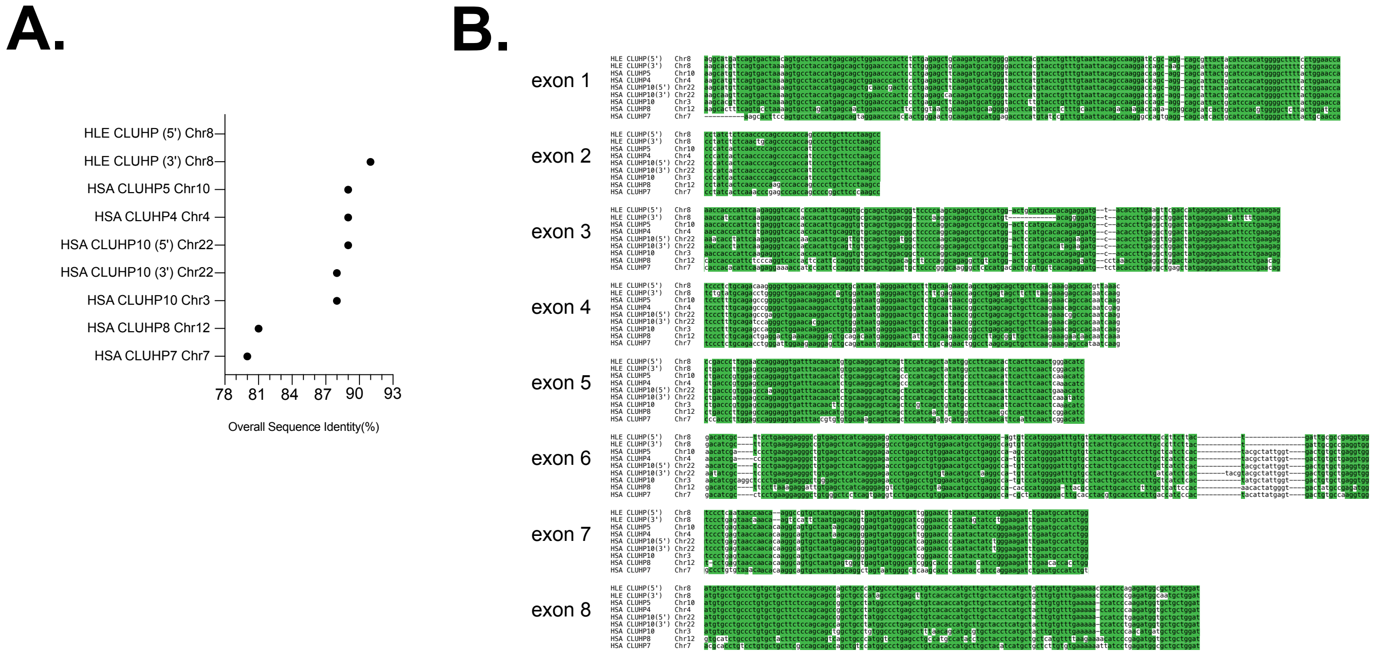

### Hartley et al, Figure S8

A.

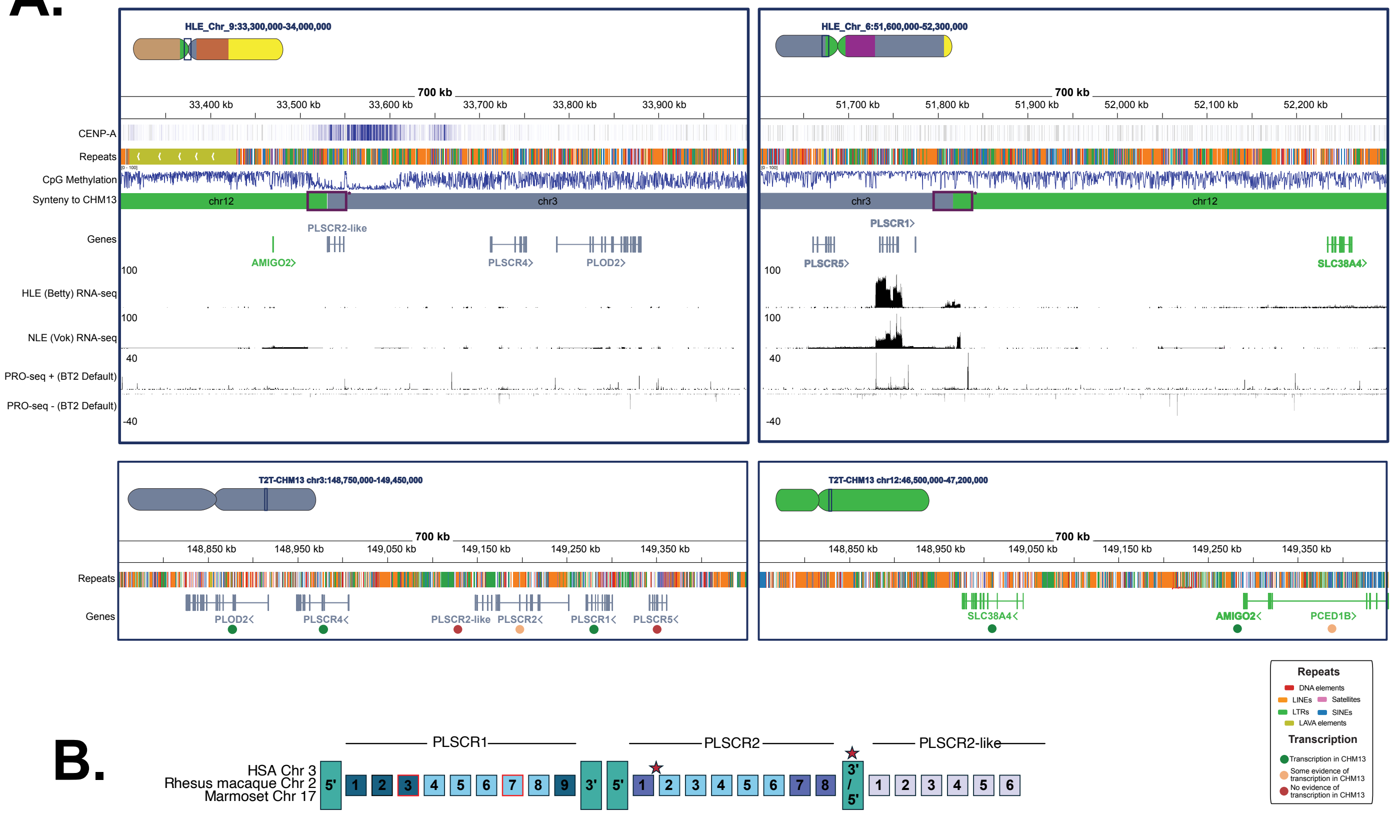


B.

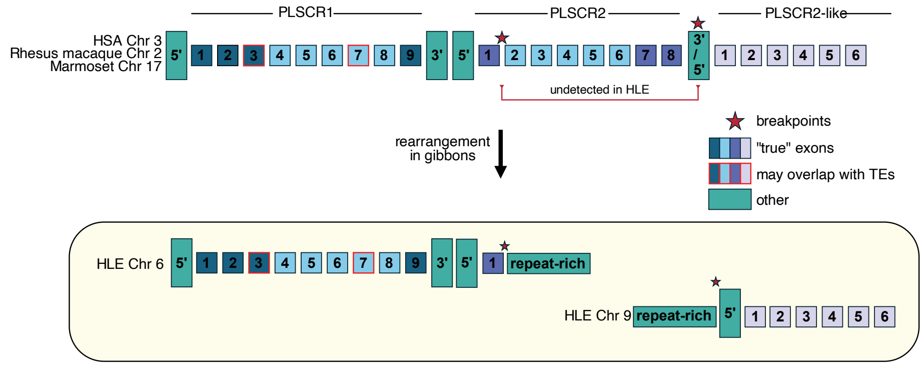

**Hartley, et al Fig S9**

A.

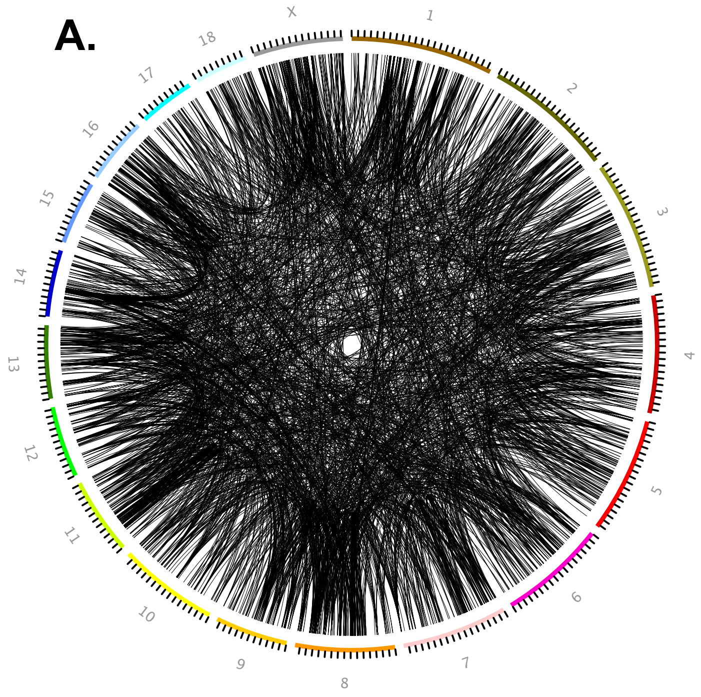


B.

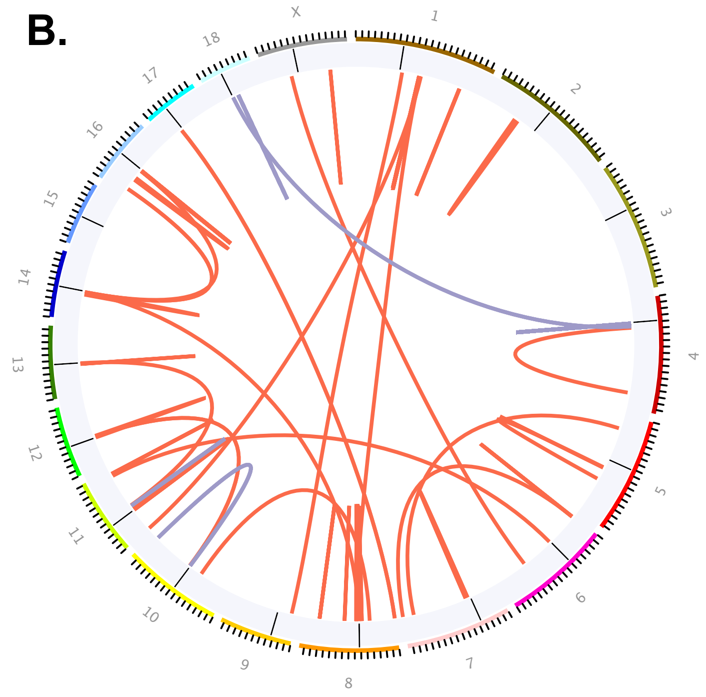

### Hartley et al, Figure S10

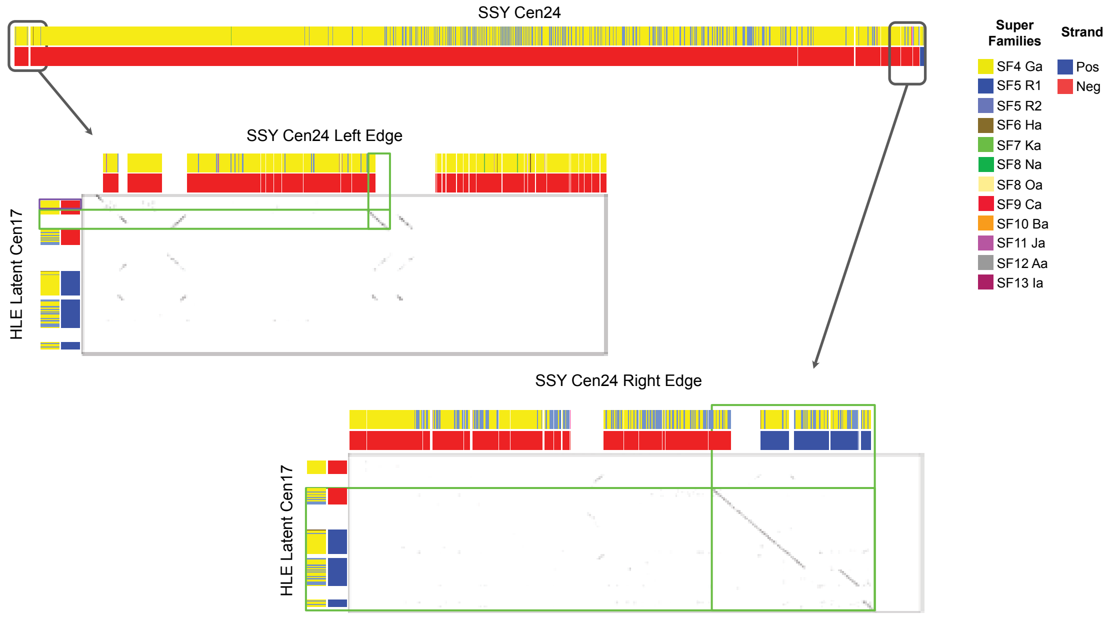

### Hartley et al, Figure S11

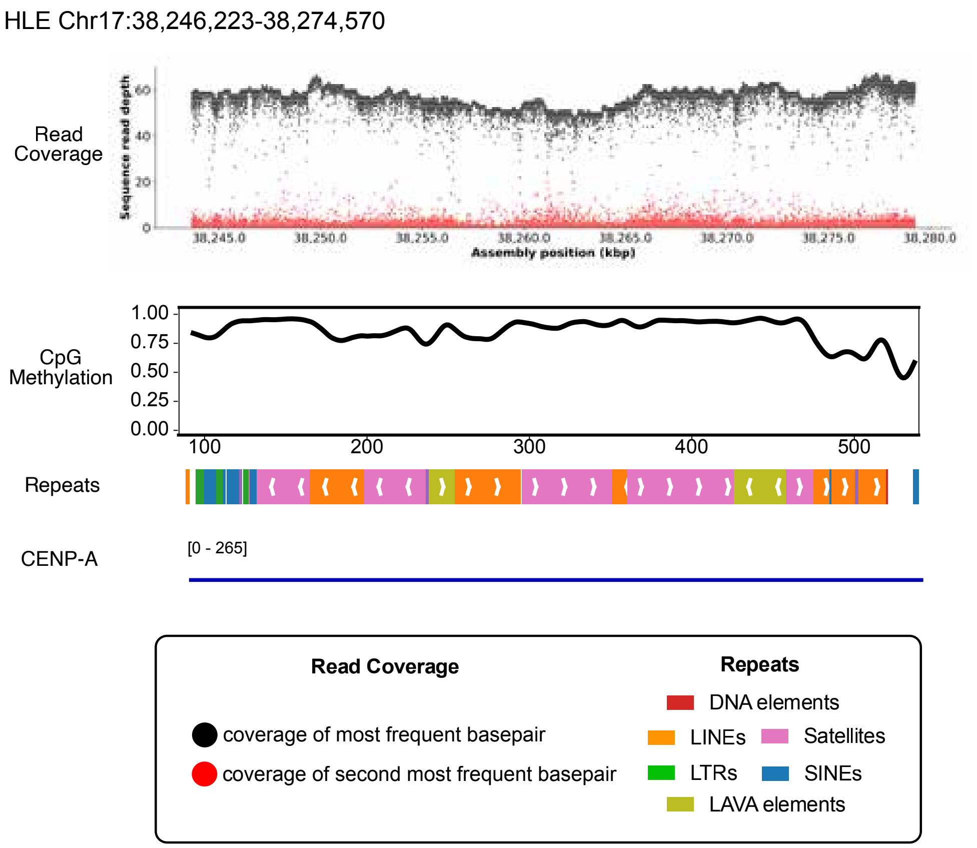

#### Hartley et al, Figure S12

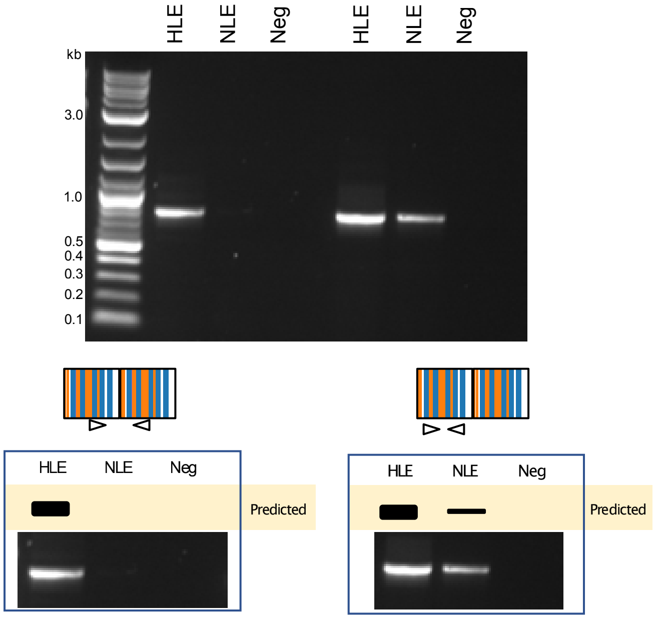

### Hartley, et al _ Figure S13

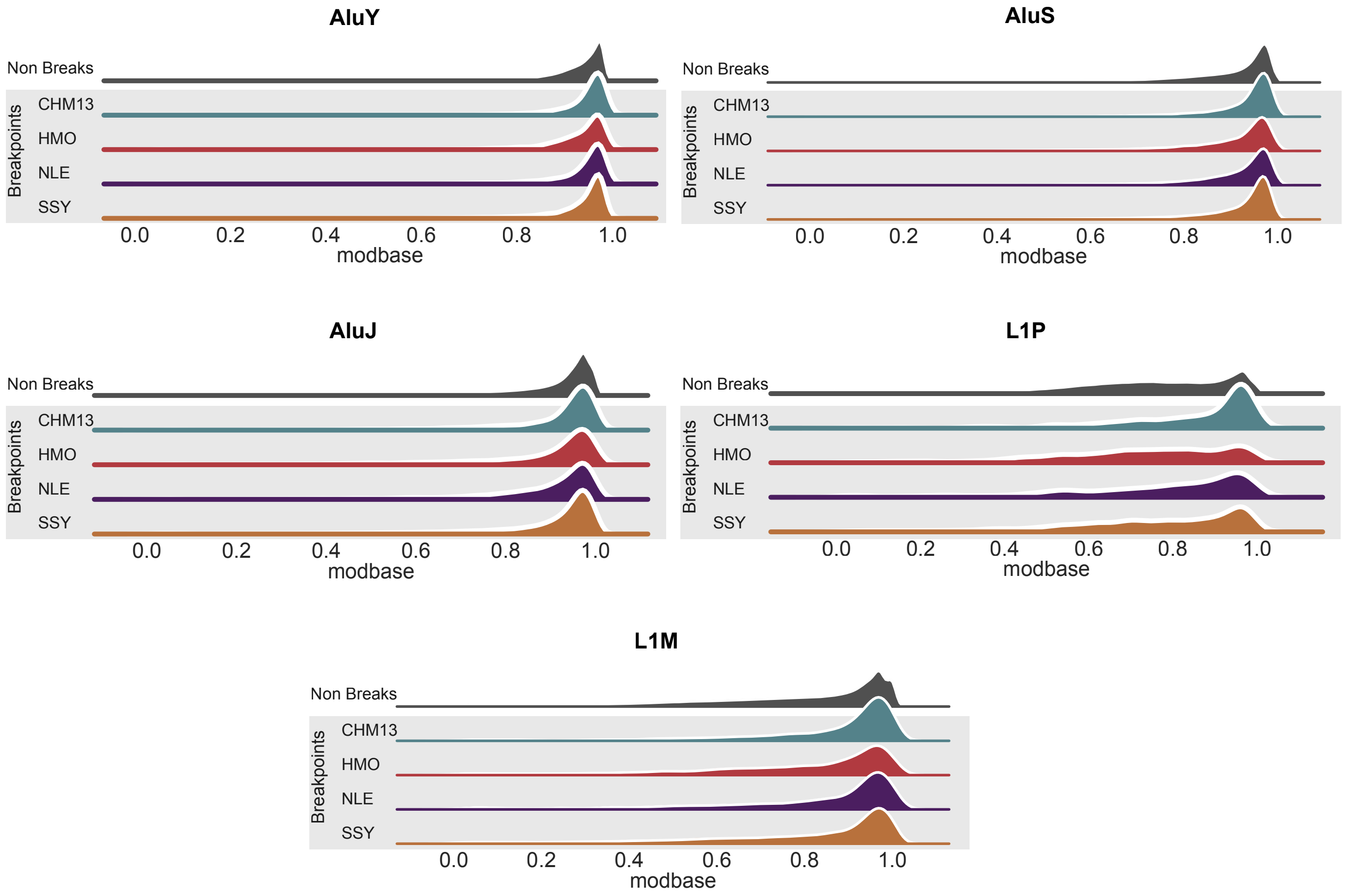

Hartley, et al _ Figure S14

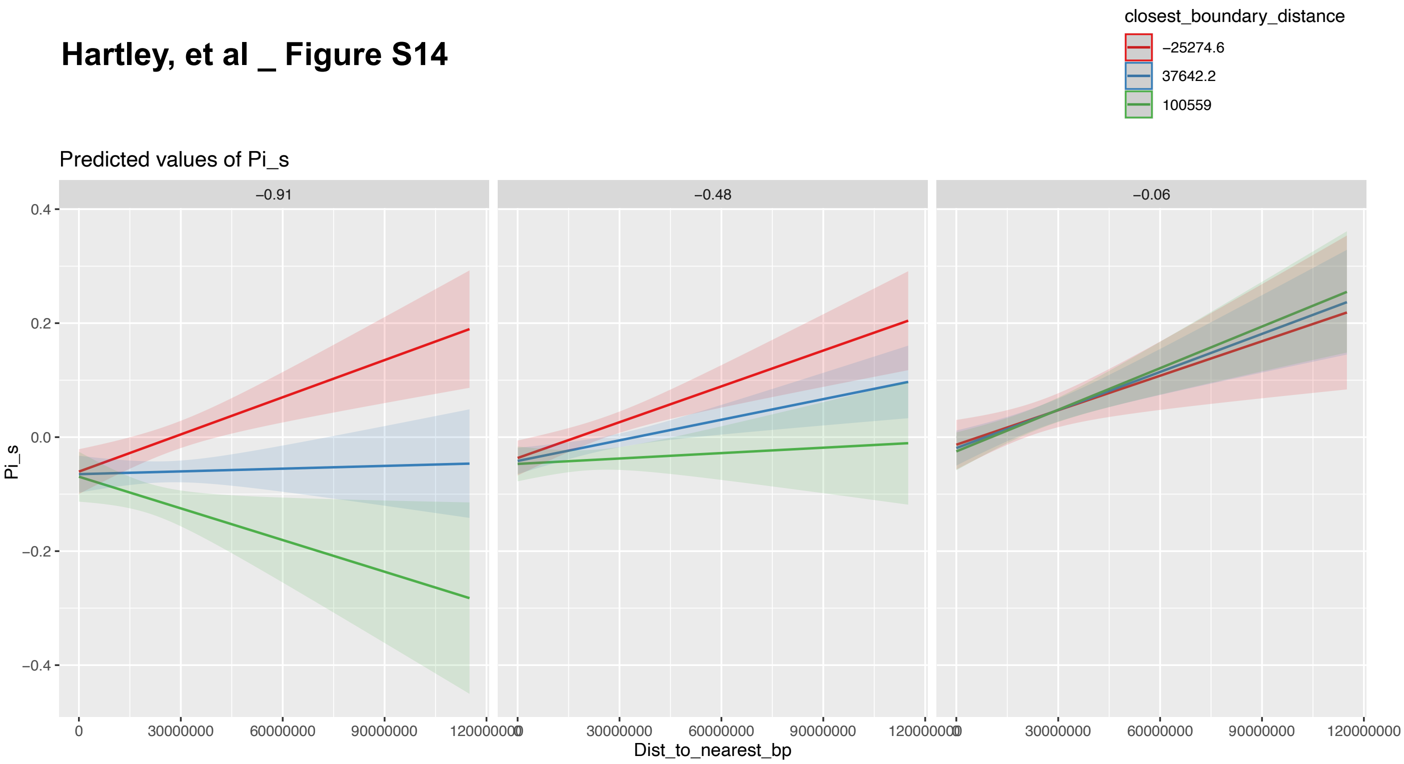

**Hartley, et al _ Figure S15**

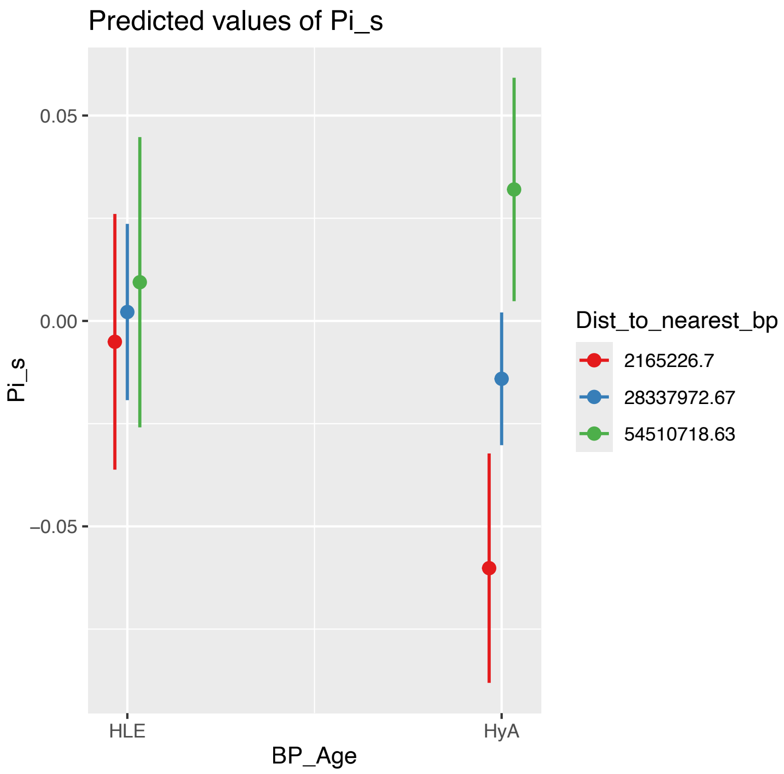

### Hartley, et al _ Figure S16

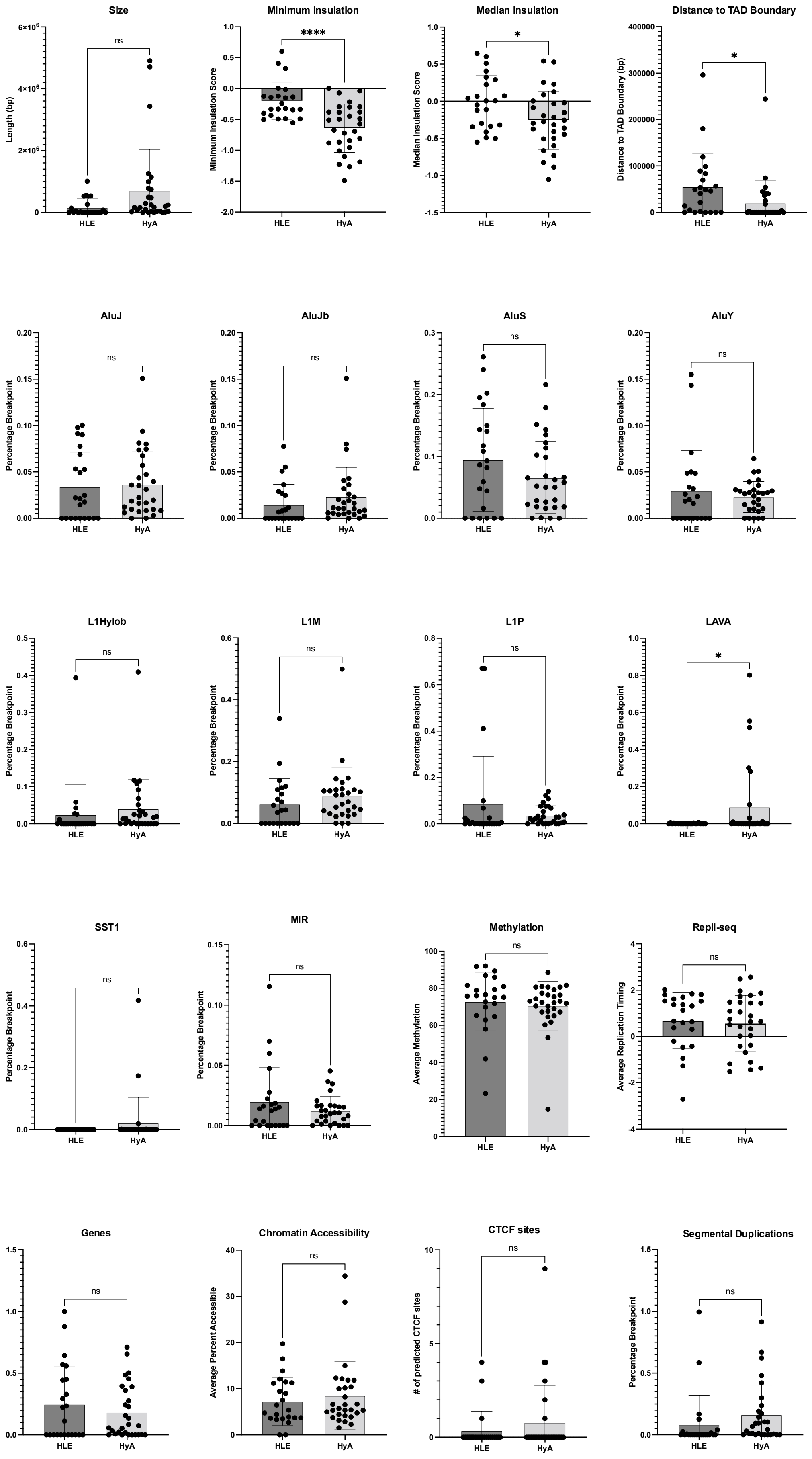
